# Amyloid forming human lysozyme intermediates are stabilised by non-native amide-π interactions

**DOI:** 10.1101/2024.12.20.629167

**Authors:** Minkoo Ahn, Julian O. Streit, Christopher A. Waudby, Tomasz Włodarski, Angelo Miguel Figueiredo, John Christodoulou, Janet R. Kumita

## Abstract

Mutational variants of human lysozyme cause a rare but fatal hereditary form of systemic amyloidosis by populating an intermediate state that self-assembles into amyloid fibrils. Despite its significance in lysozyme amyloidosis, the intermediate state has been recalcitrant to detailed structural investigation as it is only transiently and sparsely populated. Here, we investigated the intermediate state of a mutational variant of human lysozyme (I59T) using CEST and CPMG RD NMR at low pH. ^15^N CEST profiles probed the thermal unfolding of the native state into the denatured ensemble and revealed an additional state distinct from the two major states. Global fitting of ^15^N CEST and CPMG data provided kinetic and thermodynamic parameters for the exchange between all three states, characterising the intermediate state populated at 0.6%. ^1^H CEST data also confirmed the presence of the intermediate state displaying unusually high or low ^1^H_N_ chemical shifts. To further investigate the structural details of the intermediate state we used molecular dynamics (MD) simulations, which recapitulated the experimentally observed folding pathway and free energy landscape. A high-energy intermediate state with a locally disordered β-domain and C-helix was observed, revealing non-native hydrogen bonding and amide-π interactions. These interactions account for the anomalous ^1^H chemical shifts and likely stabilise the transient intermediate state structure. Together, our NMR and MD data provide the first direct structural information on the intermediate state, offering insights into targeting lysozyme amyloidosis.

## Introduction

Many globular proteins undergo locally cooperative unfolding, forming transient intermediate states that self-assemble into amyloid fibrils associated with neurodegenerative and systemic diseases^1,2^. Human lysozyme is a globular protein with a persistent native structure, but mutations in its genetic code cause it to populate a partially unfolded intermediate state, which aggregates and accumulates in the viscera of patients, reaching up to kilogram quantities^3^. It is thought that the intermediate state retains the native β-domain structure, while the β-domain is mostly denatured^4-6^. The β-sheet region of the β-domain is of particular importance, as disease-related mutations occur in this region^3^, it is selectively denatured in the intermediate state^4,6^, and establishes inter-molecular interactions^7^ to form the fibril core in amyloid^8^. The disease-related variants have a higher propensity to populate the intermediate state^4,6,9^ and aggregate more rapidly than the wild-type (WT)^10^.

Previously, we investigated the local cooperativity in the thermal unfolding of the WT and mutational variants (I56T and I59T) of human lysozyme under acidic pH conditions^6^. Using 2D HSQC spectra and both far- and near-UV CD spectra with increasing temperatures, we revealed a pseudo-two-state unfolding mechanism, which involves a cooperative loss of native tertiary structure, followed by the progressive unfolding of the denatured ensemble^6^. The intermediate state, considered one of many protein conformers within the denatured ensemble, was not structurally characterised due to the lack of first-order cooperative unfolding and detectable heat in calorimetric analysis. However, recent advances in solution NMR spectroscopy, particularly chemical exchange saturation transfer (CEST) and CPMG relaxation dispersion (RD) NMR, have enabled the observation of protein dynamics across a broad range of timescales, providing insights into the exchange processes between protein conformers at millisecond to microsecond timescales^11^.

To understand the unfolding process of human lysozyme and characterise the involved protein conformers, we harnessed ^15^N CEST NMR at pH 1.2 to monitor the unfolding of the I59T variant, which shares key attributes with disease-associated variants^12. 15^N CEST detects the exchange between the native state and the denatured ensemble, particularly revealing the latter at low temperatures, where peaks are invisible in 2D HSQC spectra. In addition, a distinct minor state was observed by both ^15^N and ^1^H (H_N_, H_α_, H_γ_ and H_δ_) CEST experiments. This intermediate state was observed only in the N-terminus, β-domain and C-helix, which unfold at lower temperatures compared to other regions of the protein (α-domain). By fitting all ^15^N CEST and CPMG data into a three-state unfolding model, we determined the exchange parameters and population of all three states, showing that the intermediate state is marginally populated (∼0.6%) near the melting temperature (∼35 ºC). The H_N_ CEST data confirmed the same exchange process and revealed unusually high or low chemical shift values for the intermediate state.

We further investigated the structural features of the intermediate state using ratchet-and-pawl molecular dynamics (MD)^13^ simulations to explore the folding free energy landscape. Our simulations from two different force fields accurately and independently recapitulate the folding process of lysozyme and reveal a high-energy local minimum consistent with a transient equilibrium folding intermediate, which has an overall stable compact structure with a locally disordered β-domain and C-helix. Extensive unbiased simulations of these intermediate state structures reveal the molecular basis of the anomalous ^1^H_N_ chemical shifts from the CEST data: structures with non-native hydrogen bonding and amide-π interactions are observed, and these are the likely cause for the strong deshielding and shielding effects, respectively, on the observed proton nuclei.

Collectively, our NMR and MD simulations enable the first direct observation of the lowly populated lysozyme intermediate state and reveal how these disease-causing structures are stabilised by non-native interactions, providing a potential structural target for perturbing lysozyme fibril formation in systemic amyloidosis.

## Results

### The invisible denatured ensemble observed by ^15^N CEST during thermal unfolding

The thermal unfolding of I59T at pH 1.2 was previously monitored by recording ^1^H-^15^N HSQC spectra and revealed a pseudo-two state unfolding process where the native state (N) exchanges with the denatured ensemble (D), which progressively unfolds its secondary structure as the temperature increases^6^. I59T predominantly populates the N state at 25 ºC and the D state at 45 ºC, while at 35 ºC, close to its melting temperature (T_m_) for tertiary structure (determined by near-UV CD spectroscopy, Fig.1e)^6^, both N and D peaks are observed (Fig.1a, S1). As the temperature increases, different regions of the protein gradually show denatured peaks, with the N-terminal short β-strand (residues 2-4) and the three β-strands in the β-domain (residues 40-55) predominantly unfolding at low temperatures (< 30 ºC, Fig.1b-c, e).

To further investigate the equilibrium between the native and denatured states of lysozyme during thermal unfolding, we performed ^15^N CEST NMR experiments, which probe exchange on the second to millisecond timescales^14^. Fig.1d shows the CEST profiles for two residues (V2 and L25) at 30 ºC. V2, which displays its D peak at the lowest temperature (from 5 ºC), shows clear N-D state exchange at 30 ºC (Fig. 1d). Similarly, L25, located in one of the α-helices (helix B), shows N-D exchange at 30 ºC, even though its D peak is only visible at higher temperatures (> 35 ºC) in the HSQC spectra. At 25 ºC, all visible N state peaks (109 out of 118) in the 2D spectra exhibit exchange with the D state in the CEST profile (Fig.1b), except for residues with the same chemical shifts for both N and D states, preventing direct detection of exchange. This suggests that even at low temperatures, the invisible protein conformers of the D state, which are in equilibrium with the N state, can be observed using CEST, enabling us to explore the detailed exchange processes during thermal unfolding.

**Figure 1.**
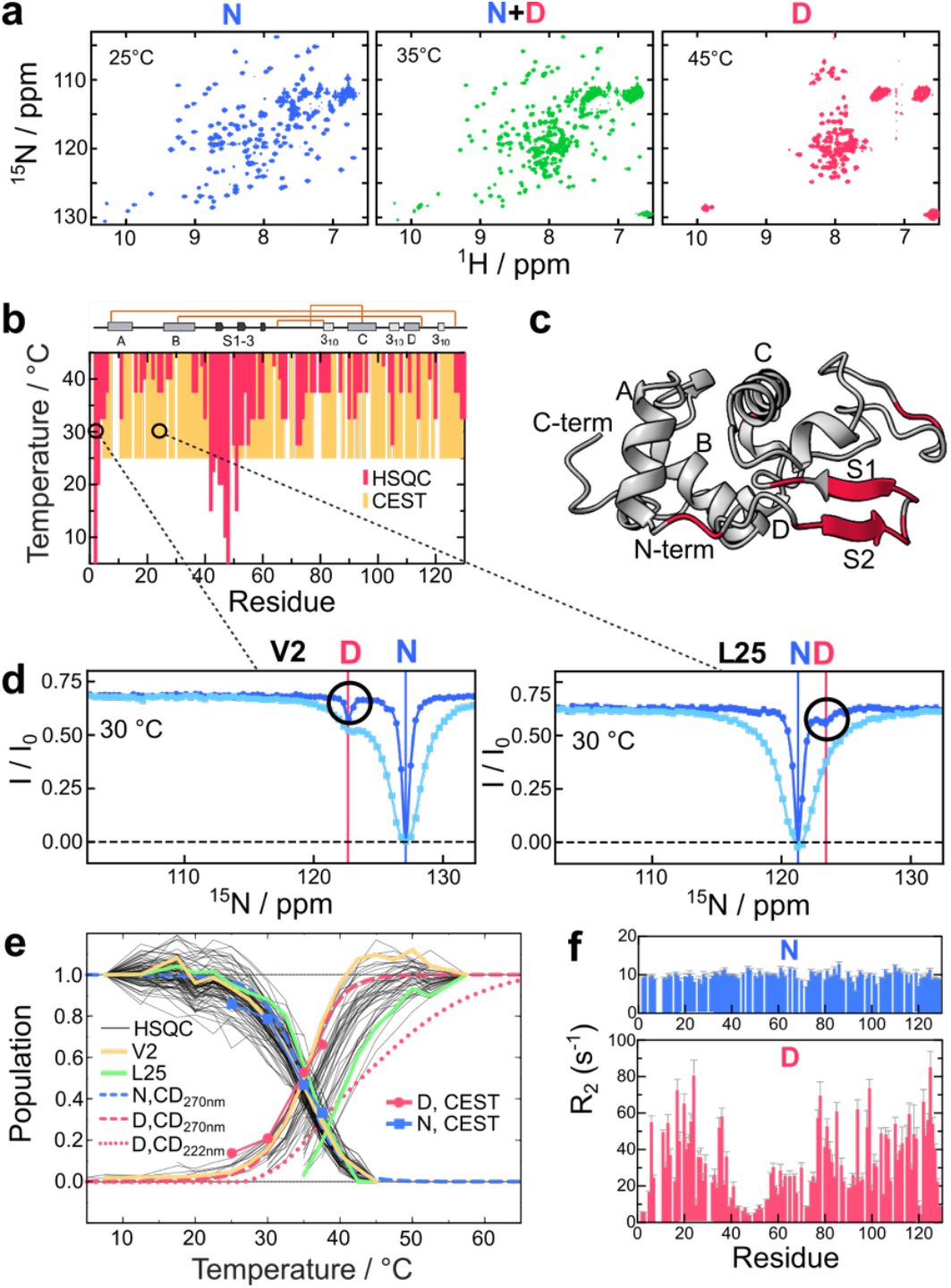
The denatured ensemble state observed by CEST NMR. (a) ^1^H-^15^N HSQC spectra of I59T at pH 1.2 at three different temperatures (25, 35 and 45 ºC). (b) Observation of the denatured state using ^1^H-^15^N HSQC (5-45 ºC, red) and ^15^N CEST NMR (25-45 ºC, yellow). (c) Residues showing denatured peaks below 30 ºC, highlighted on the I59T structure (PDB ID:2meh)^16^. (d) ^15^N CEST profile for V2 and L25 at 30 ºC. (e) Thermal unfolding of I59T monitored by NMR and CD spectroscopy. Peak volumes in the HSQC spectra are shown in black solid lines, with data from V2 and L25 in yellow and green, respectively. Thermal unfolding by near-UV CD (270nm) was fit to a two-state unfolding model, with native (N, blue dashed) and denatured ensemble (D, red dashed) populations indicated. Far-UV CD (222nm) thermal unfolding data were normalised, and the denatured population is shown as a red dotted line. ^15^N CEST data from all observed peaks (109 residues) were globally fit to a two-state unfolding model, with N (blue squares with blue solid line) and D (red circles with red solid line) populations shown. (f) ^15^N transverse relaxation rates (R_2_) of the N (blue) and D (red) states derived from ^15^N CEST fitting at 35 ºC.

The CEST profiles of 109 residues were globally fit to a two-state model, revealing the populations of both the N and D states consistent with near-UV CD results (Fig.1e)^6^. Fig.1f shows the R_2_ values of the N and D states derived from CEST data fitting. The N state exhibits uniform R_2_ values of ∼10 s^-1^ across the protein sequence, indicating a stable protein structure. In contrast, the D state displays variable R_2_ values: residues that unfold at lower temperatures (N-terminal residues 2-4 and β-strands residues 40-55) have lower R_2_ values (< 10 s^-1^), indicating fast dynamics. The rest of the protein, particularly the α-domain, shows higher R_2_ values, reflecting additional rapid exchange processes among protein conformers^15^ with varying degrees of α-domain denaturation. The ^15^N R_2_ values of the D state follow a similar trend to experimentally measured ^1^H_N_ R_2_ values^7^, indicating consistent D state dynamics under these conditions and explaining the lower D state populations observed in the α-domain peaks of the HSQC spectra, as exemplified by L25 (Fig.1e).

### The invisible intermediate state observed by ^15^N CEST and CPMG RD

To quantitatively investigate the thermal unfolding and associated exchange processes, we analysed the ^15^N CEST data collected at 25-45 ºC, which encompass the melting temperatures of both tertiary (35 ºC) and secondary (45 ºC) structures^6^. Fig. 2a shows the CEST profiles of A42, a residue on the S1 strand of the β-domain. The CEST profiles from the N and D peaks recorded at two different B1 fields (15 and 60 Hz) reveal a temperature-dependent increase in the D state population and a corresponding decrease in the N state population, shown in the changing depth of the minor state CEST dips. Additionally, a distinct CEST dip is observed at a ^15^N shift of 131.4 ppm, indicating a third state in exchange with both the N and D states. This additional state, not an artifact from H/D exchange^17^, was observed in the CEST profiles of 14 residues, located in the β-domain (A42, T43, S51, T52, D53, G55, I56, F57, Q58, T59, N60, D67, L84) and the C-helix (A92) (Fig.2c-d, S2). These β-domain and the C-helix residues are known to be locally denatured in the intermediate state populated during unfolding^4,5^ and form the core of amyloid fibrils^8^. Moreover, most disease-related point mutations occur in this region^3^. These findings suggest that the third state uniquely observed in the CEST profiles is likely to be the unfolding intermediate (I) state that self-assembles into amyloid fibrils. Additionally, V2 at the N-terminus, which populates its D state at a very low temperature like the β-strand residues (Fig.1b), also shows the I state, indicating the role of the N-terminal short β-strand (residue 2-4) in N state stability and unfolding^18^. The I state is observed from all CEST profiles recorded at different field strengths (500, 700 and 950 MHz) probing consistent exchange processes (Fig. S3).

**Figure 2.**
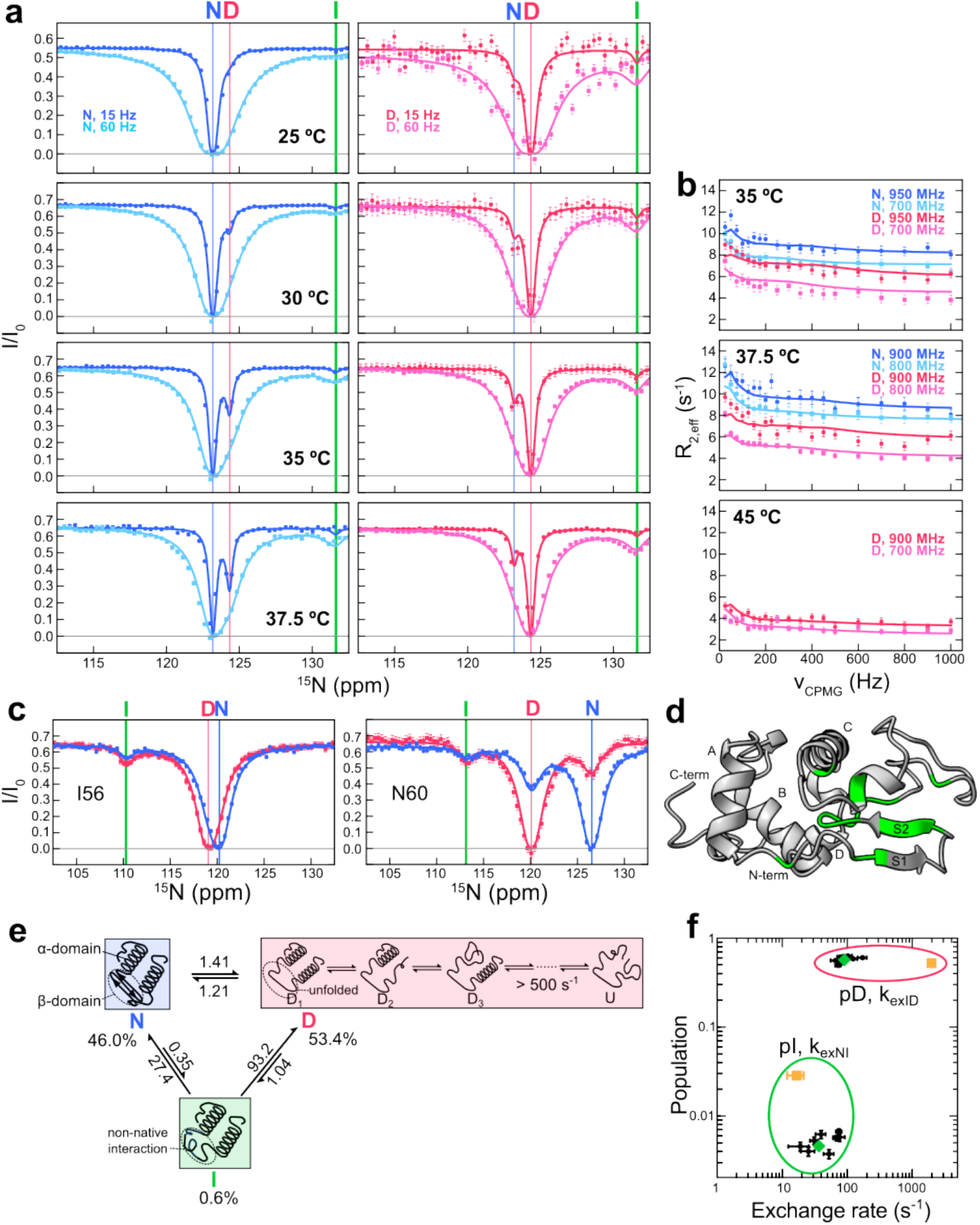
The intermediate state observed by ^15^N CEST and CPMG RD NMR. (a) ^15^N CEST and (b) CPMG RD data for A42 during thermal unfolding (25-45ºC). The chemical shifts of the native (N), denatured ensemble (D) and intermediate (I) states are shown in blue, red and green vertical lines, respectively. (c) CEST profiles of I56 and N60 at B_1_= 60Hz. The presence of the I state is indicated by an additional CEST dip, marked by green vertical lines. (d) Residues showing the I state CEST dips distinct from the D state, highlighted on the I59T structure (PDB ID:2meh)^16^ (e) A three-state thermal unfolding model used for the global fitting of the CEST and CPMG data from 14 residues in (d). The kinetic parameters are derived from data at 35 ºC. N, I and D represent the native, intermediate and denatured states based on the β-domain and C-helix structures. D_1_, D_2_, D_3_ and U represent rapidly exchanging protein conformers that make up the D state, with the U indicating a fully unfolded protein. (f) Comparison of populations and exchange rates from the global fitting of the ^15^N CEST and CPMG RD data from individual β-domain residues (black dots), all β-domain and C-helix residues together (green diamond) and α-helix (helix B) residues (orange square). Data points in the red and green ellipses represent the population of the D state (with K_ex_ID_) and the I state (with K_ex_NI_), respectively.

To further characterise the exchange processes observed by CEST, we performed ^15^N CPMG RD experiments at multiple field strengths (700, 800, 900 and 950 MHz), which probe faster millisecond exchanges^11^ (Fig.2b, S3). The CPMG RD data from both the N and D peaks of the 14 residues at the N-terminus, β-domain and C-helix show exchange processes that were quenched at high CPMG frequencies (v_CPMG_), revealing the presence of the additional state(s) in millisecond exchange with the N and D states (Fig. 2b, S3). We globally fitted all CEST and CPMG data from various temperatures and field strengths for each residue showing distinct I state in the CEST profiles to three-state sequential exchange models (N ↔ I ↔ D, N ↔ D ↔ I and I ↔ N ↔ D), as well as a triangular model that includes all possible exchange processes (Fig.2e, S4). The triangular model provided the best fit with the lowest reduced χ^2^, BIC and AIC values (Fig. S4), suggesting that the I state is not an off pathway intermediate in the unfolding/folding process. Fitting the data for each β-domain residue resulted in consistent populations and exchange rates, and the global fitting of all these residues together yielded the same kinetic parameters (Fig.2f). These results suggest that the I state is populated through the local cooperative unfolding of the β-domain and the C-helix, only transiently and marginally during the thermal unfolding process with a maximum population of 0.6 % (35 - 37.5 ºC), which is near the lower detection limit of both CEST and CPMG RD experiments^11^. The exchange rates (k_ex_) between N and I, and I and D are 27.7 and 94.3 s^-1^ at 35 ºC, with free energy differences (ΔΔG) of 2.7 and -3.06 kcal/mol, respectively.

Unlike the β-domain and C-helix residues, the rest of the α-helix residues in the α-domain show no indication of the I state in their CEST profiles, with D state peaks only appearing at higher temperatures (Fig.1b). The progressive thermal unfolding of α-helices leads to the formation of a rapidly interconverting ensemble of protein conformers with varying degrees of denaturation^6^. As a result, α-domain residues exhibit distinct CEST and CPMG profiles compared to the β-domain and C-helix residues. Their D state minor CEST dips are broader than that of the β-domain and C-helix residues (Fig. 1f, S5a) and their N state CPMG peaks show smaller R_ex_ values (< 2 s^-1^, Fig. S5b), both indicating faster exchange between interconverting protein conformers^14^. The absence of I state CEST dips and the limited number of D state peaks for α-domain residues make fitting their data into a three-state unfolding model challenging, resulting in noticeably different kinetic parameters (Fig.2f). We then attempted to fit the α-domain data with the β-domain data into the three-state model using either the N or D state chemical shifts for the I state. The results showed better fitting when the I state chemical shifts were assigned to the N state, suggesting that the I state retains a native-like α-domain structure, consistent with the predicted I state structure^4^. The R_2,0_ values from the CPMG analysis of denatured peaks suggest the presence of faster exchange in α-domain residues (Fig. S5c). We recorded rotating frame (R_1π_) relaxation dispersion (RD) for an α-domain residue at the C-terminus (G129), and the global fitting of both on-/off-resonance R_1π_ RD data reveal a faster exchange process between these protein conformers (Fig. S5d). These NMR experiments collectively reveal fast (> 500 s^-1^) exchange processes between protein conformers with varying degrees of α-domain denaturation (Fig. 2e).

As previously reported, I59T exhibits reduced global cooperativity of unfolding in comparison to the WT protein, making this variant more aggregation-prone^10^. We performed CEST experiments on the WT protein at multiple temperatures (25, 35 and 45 ºC) (Fig. S6), and found no evidence of the I state. This is likely due to the lower population of the I state in the WT^4,10^, which falls below the detection limit of CEST experiments.

### ^1^H chemical shifts of the intermediate state

To further investigate the intermediate (I) state of I59T observed in ^15^N CEST and CPMG experiments, we performed CEST experiments on other NMR-observable nuclei including ^13^C and ^1^H. Both backbone (H_N_, C_α_/H_α_) and side chain nuclei (C_γ_/H_γ_, and C_δ_/H_δ_) CEST data revealed exchange between the N and D states, detecting the invisible D state peaks at lower temperatures (Fig. S7) like the ^15^N CEST results (Fig.1b). However, no intermediate state was observed in the ^13^C CEST, potentially due to the inherently low sensitivity of methyl CEST^14^.

In contrast, the ^1^H CEST data revealed both invisible D and I states. Figure 3a shows the HSQC spectrum of I59T at 35 ºC with ^15^N and ^1^H CEST profiles of three β-domain residues (A42, I56 and T59). The anti-phase absorptive ^1^H_N_ CEST profiles clearly show the I state in these β-domain residues in exchange with both N and D states (Fig.3a). Interestingly, the I state exhibits significant chemical shift differences from both the N and D states in the ^1^H and ^15^N CEST data, indicating strong shielding (I56, I59) or deshielding (A42) effects on the observed nuclei. The I state was also detected in the ^1^H CEST profiles of H_α_ and H _δ_nuclei of A42 and I56 (Fig.3b-c), further confirming its presence.

**Figure 3.**
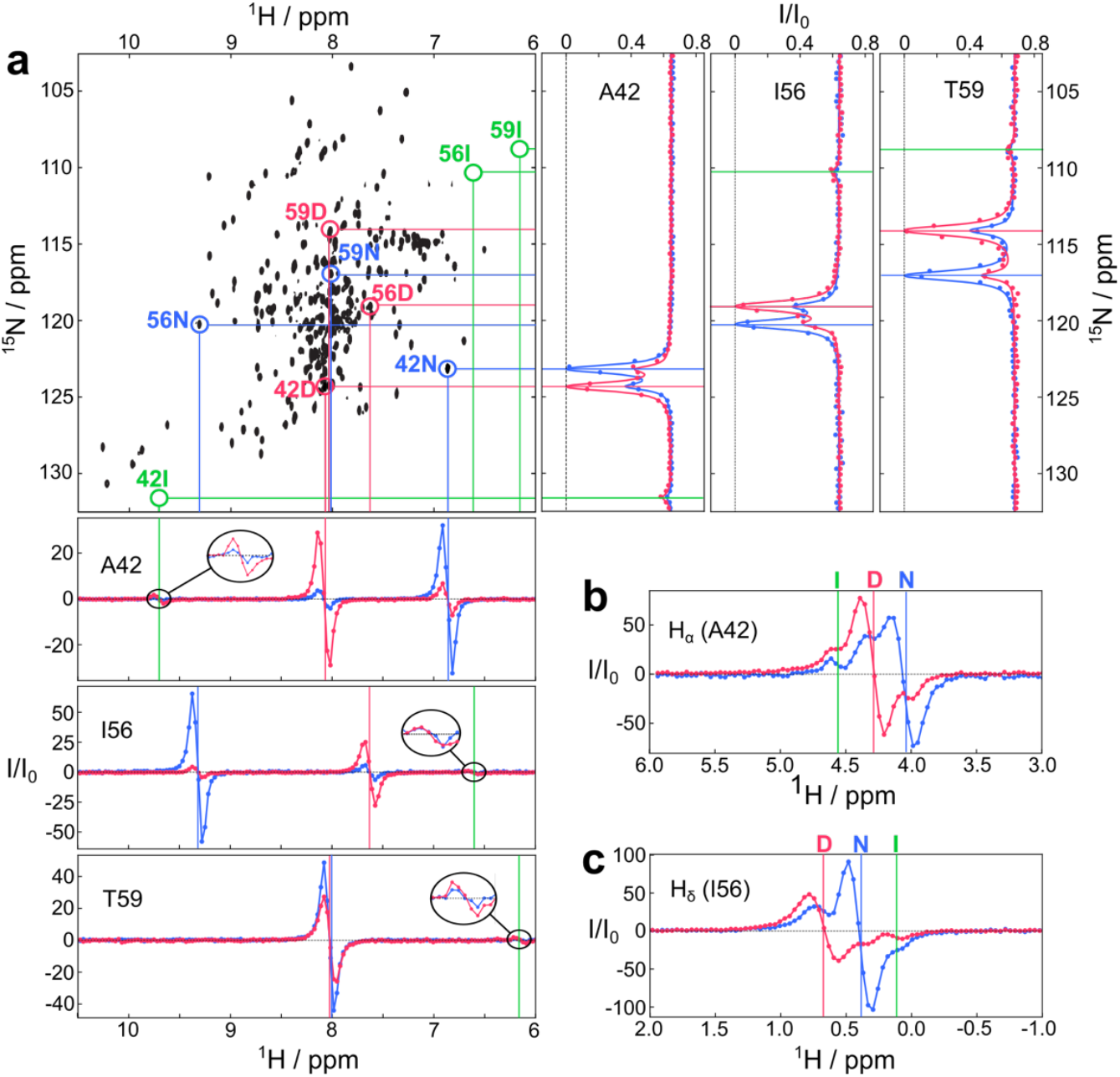
The I state monitored by ^1^H CEST. (a) ^1^H_N_ and ^15^N CEST data for A42, I56 and T59 shown with the ^1^H-^15^N HSQC spectrum at 35 ºC. ^1^H_N_ CEST were recorded at a ^1^H frequency of 950 MHz. Blue and red dots connected by lines represent the native (N) and denatured (D) peaks, respectively. Green circles indicate the I state in exchange with both N and D. (b) H_α_ CEST of A42 and (c) H_δ_ CEST of I56 both revealing the presence of the I state.

### MD simulations reveal a compact intermediate state with unstructured β-domain and C-helix

Next, we aimed to investigate the structure of the intermediate state and sampled the folding free energy landscape of the I59T using all-atom MD simulations in explicit solvent. Protein folding simulations are challenging due to the long timescales involved and complex folding free energy landscape, thus we took the advantage of an enhanced sampling approach referred to as adiabatic MD or ratchet-and-pawl MD simulations (rMD, see methods)^13,19-22^. In this method, we apply a biasing force when the protein backtracks along a pre-defined collective variable that captures folding progression (defined by i.e., the number of established native contacts). This approach is computationally efficient and has been successfully applied to various proteins in implicit and explicit solvent^13,19-22^. Starting from 50 completely unfolded structures (with a fraction of native contacts below 0.1) with all four disulfide bonds oxidised (confirmed to be intact by the Cβ chemical shift of the Cys residues, Table S1), we generated a total of 1,800 trial folding trajectories (50×36) for two different force fields, C36m^23^ and a99sb-disp^24^, of which 415 and 368 reached the native state in 5 ns of biased simulation time, respectively. We then calculated the kinetic folding free energy landscape as a function of two collective variables, the Cα RMSD to the native state and Q (the fraction of native contacts) for each force field (Fig. 4a, Fig.S8a for C36m). These energy landscapes contain a deep free energy minimum corresponding to the native state, and additionally a high-energy local minimum consistent with a transient, on-pathway kinetic folding intermediate. To identify the folding intermediate structures, we clustered conformations obtained by rMD, which belong to the intermediate state energy minimum. The most populated cluster (a99sb-disp) represents a state with an unfolded β-domain and a partially unfolded C-helix (Fig. 4b), consistent with previous experimental studies^6^ and our ^15^N CEST experiments. A similar state is observed when clustering the intermediate state region obtained with the C36m force field (Fig. S8a-b).

**Figure 4.**
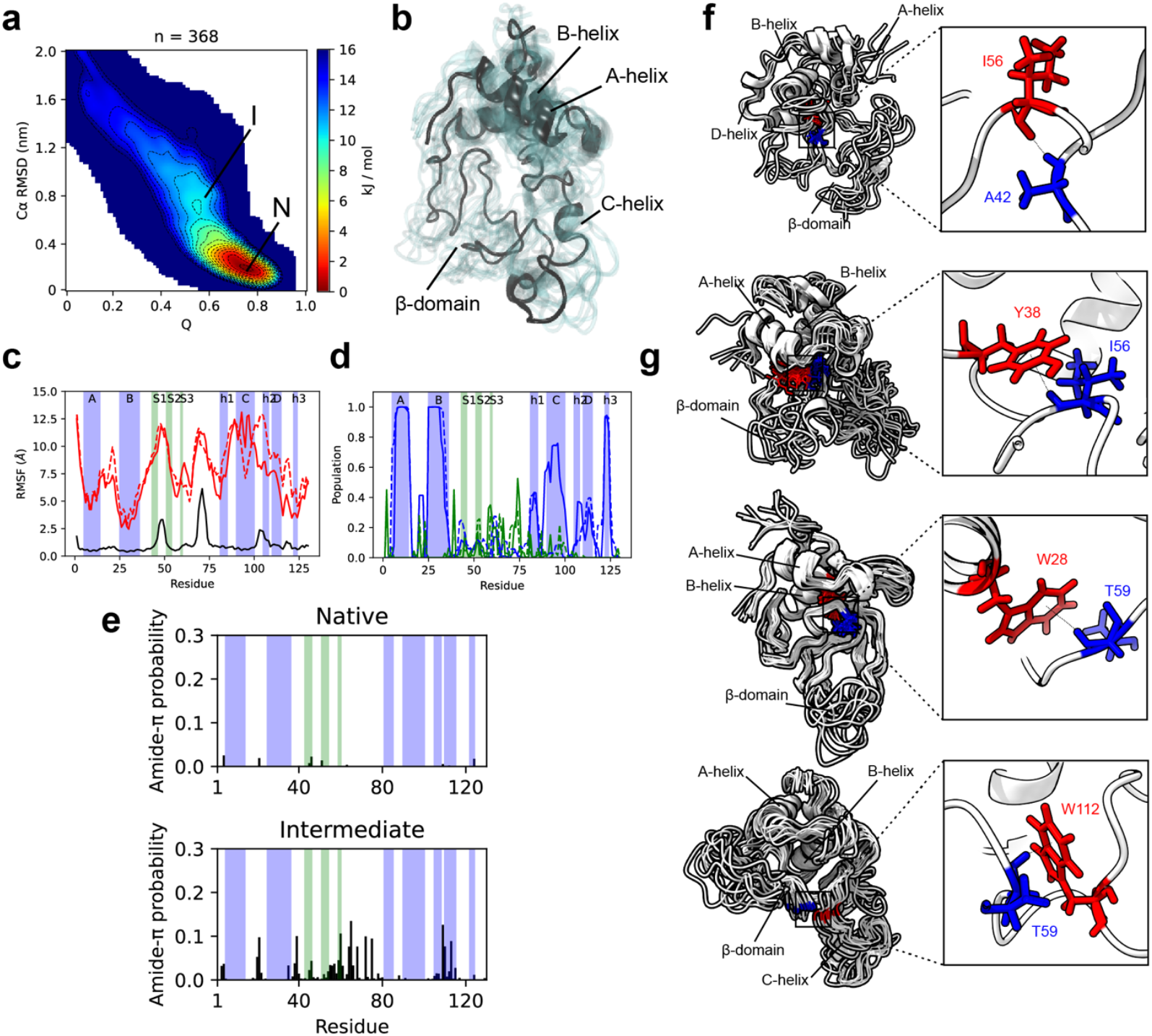
Characterisation of the kinetic folding free energy landscape and intermediate state by molecular dynamics simulations. (a) Kinetic folding free energy landscape of I59T calculated from all (n = 368) rMD trajectories that reach the native state with the a99sb-disp force field. The regions of the native state and metastable intermediate are annotated. (b) Representative ensemble of the folding intermediate state (I) from panel a. This ensemble represents the most populated cluster from the I state region in panel a. (c) Backbone dynamics of the native (black) and intermediate state (red) calculated from the unbiased MD simulations for the a99sb-disp force field, quantified by the RMSF (Cα atoms). The solid and dashed red lines represent the RMSF calculated from two halves of the ensemble (each from 4×10μs of unbiased MD simulations for a different intermediate starting structure to assess the reproducibility of the structural ensembles). (d) Average secondary structure populations (α-helix – blue, β-sheet – green) of the intermediate state ensemble obtained with a99sb-disp (solid and dashed lines represent the two half ensembles). (e) Ensemble-averaged probability of forming amide-π interactions in the native and intermediate state ensemble obtained with a99sb-disp. (f)-(g) Non-native interactions observed in the β-domain rationalising the intermediate state chemical shifts. (f) The amide of residue A42 donates both mainchain and sidechain hydrogen bonds. The ensemble and interaction shown highlight a backbone hydrogen bond donated to I56. (g) The I56 amide group participates in amide-π hydrogen bonding and an example sub-ensemble and interaction with Y38 are shown. The amide bond is oriented perpendicular to the ring plane. The T59 amide group also participates in amide-π hydrogen bonding. An example ensemble and interaction with W28 are shown (sampled in a C36m simulation), as well as with W112 (sampled in an a99sb-disp simulation).

The average folding pathway observed in the rMD simulations qualitatively recapitulates the observed folding pathway from biophysical experiments (SI text), demonstrating the delayed folding of β-sheet and C-helix in comparison to the fast-folding of remaining α-helices (Fig. S8c-e, SI text). This is consistent with the intermediate state obtained from clustering, which lacks native β-sheet and C-helix structures. An analysis of the solvent accessibility of the indole amide (W109 and W112) and side chains of Trp residues (W28, W64, W109 and W112) also describe the same folding pathway (Fig. S8f-h) observed by stopped-flow CD and NMR hydrogen-deuterium exchange experiments^25^, demonstrating the delayed folding of the C-helix (SI text). Overall, our rMD simulations predict the folding pathway from experimental studies with helices A-C forming secondary structure rapidly, followed by helix D and then the β-sheet. All structural elements except the C-helix form tertiary contacts early, while the C-helix folds late against the β-sheet and preformed α-domain. The intermediate state is also consistent with the I59T mutation as this increases its population by destabilising the native state, since the sidechain of residue 59 is located between the β-sheet and helices h2/C.

Two representative structures from the intermediate clusters with an unfolded C-helix and β-sheet were selected and subjected to long-timescale unbiased MD simulations to explore the conformational space of the intermediate state further and assess its stability. We obtained 4×10μs of MD data per input structure (160μs total across both force fields) and ran unbiased simulations of the native state as an additional control (4×2.5μs per force field, 20μs total). The native state remained highly stable with average C α-RMSDs to the crystal structure of 1.88 ± 0.28 Å, while the intermediate state remained compact with only small increases in the radius of gyration (Fig. S9a). Most of the α-domain (A, B and h3 helices) showed native-like stable structures, while the β-domain and C-helix, exhibit much higher RMSF values, indicative of their increased structural dynamics in the intermediate state (Fig 4c, Fig. S9b). Likewise, the N-terminal five residues also show a large increase in RMSF, consistent with NMR relaxation measurements indicating increased dynamics of the N-terminus^7^ and their involvement in the population of the I state in our ^15^N CEST data (Fig. 2d). Also, the analysis of secondary structure stability in the unbiased MD simulations show that the I state remained stable on the timescale of tens of microseconds in unbiased MD simulations in both force fields (Fig. 4d, Fig, S9c, SI Movies). While the intermediate state remains relatively compacted due to the lack of complete unfolding, the protein becomes significantly more exposed to the solvent relative to the native state (Fig. S9d).

Lastly, we analysed the differences in intra-chain contacts between the native and intermediate states and observed some common trends in both force fields (Fig. S9e-g). Within the β-sheet, there is a loss of native contacts while new non-native interactions appear. The β-domain also loses its native contacts with helix h1 and the N-terminus, rationalising their increased dynamics. Additionally, the β-domain forms non-native interactions with helices C and h2, consistent with the observed compactness of the intermediate state and the lack of complete unfolding. While contacts between helices A and B remain largely intact, contacts between helix A and C as well as B and h2 are lost. Helix C also loses native contacts with the nearby h1 helix, which is compensated for by non-native interactions between these structural elements. In summary, the unbiased MD simulations showed that the intermediate state identified by rMD is stable on the timescales explored here (SI Movies), consistent with the ensemble corresponding to metastable intermediate state.

### Non-native interactions stabilise the intermediate state and elicit anomalous ^**1**^**H chemical shifts**

We analysed the intermediate state structures further to understand the structural basis of the observed chemical shifts of the I state from CEST NMR experiments. Highly acidic condition for the NMR experiments (pH 1.2) poses limits for back-calculation of the chemical shifts^26^ resulting in average errors higher than that of the forward model even for the native state MD ensemble (Fig. S10a); however, both native and intermediate ensembles showed better agreement with their respective experimental dataset (Fig. S10b). Globally, the agreement with the intermediate state chemical shifts is worse compared to the native ensembles, and this appears to be dominated by the deviations in the amide proton and nitrogen chemical shifts of residues A42, I56, F57, and T59, all located in the β-domain (Fig. S10c).

To characterise the intermediate state structure even further, especially the role of the non-native interactions, we focused on residues A42, I56 and T59 that show both observable ^1^H_N_ and ^15^N chemical shifts of the intermediate state. Residue A42 has an ^1^H_N_ chemical shift of 6.90 and 9.70 ppm in the native and intermediate state, respectively, exhibiting significantly higher deshielding in the intermediate state (Fig. 3). Qualitatively, this increase in chemical shift from the native state is predicted correctly by the MD simulations, but with a smaller magnitude (Fig. S10d). Residue I56, on the other hand, shows a significantly lower chemical shift for the intermediate state (6.61 ppm) compared to its native state (9.37 ppm), indicating a strong shielding effect to the observed proton. Again, MD simulations predicted a lower chemical shift in the intermediate state with some structures showing chemical shifts of < 7 ppm as the experimental data (Fig. S10d). A similar trend is seen for T59 with a native chemical shift of 8.06 ppm and intermediate shift of 6.14 ppm, suggesting a dramatic change in the average chemical environment. Our simulations correctly predict a decrease in chemical shift but with a smaller magnitude than observed experimentally (Fig. S10d). However, predicted chemical shifts of <7 ppm are sampled by both force fields, particularly for a99sb-disp where some structures have predicted T59 proton shifts as low as ∼4 ppm. Overall, despite the large deviations from the experimental shifts, our simulations sample structures that are in qualitative agreement with the experimental data, suggesting these snapshots can provide insights into the subtle interactions sampled by these residues in the intermediate state.

We inspected the MD-derived structures with the most deshielded A42 chemical shifts (proton chemical shift > 9.5 ppm) and found that the A42 amide participates in backbone and sidechain hydrogen bonding in these structures. In simulations with the C36m force field, we found the intermediate state to predominantly donate a backbone hydrogen bond to I56 and S51 (Fig. 4f), while with a99sb-disp we mostly observed a sidechain hydrogen bond to residue N39. Thus, A42 may be involved in non-native hydrogen bonding which could rationalise its deshielded chemical shift. For residue I56, we inspected structures with a predicted proton chemical shift of less than 6 ppm. We found that these structures exclusively show the amide of I56 in close contact with aromatic rings, oriented perpendicularly to the rings. This is consistent with amide-π hydrogen bonding (Fig. 4g), which has been observed for some well-ordered proteins and can lead to extremely low amide proton chemical shifts (∼3 ppm)^27^. In the C36m simulations, the most frequent amide-π contact involved the sidechain of W28 (0.8%) and in a99sb-disp, a more frequent contact with Y38 was observed in ∼7.2% of simulation frames. These contacts are non-native and are not sampled in the native state. Lastly, T59 exhibits an even lower chemical shift in the intermediate state, and we find that simulations sample structures with a proton chemical shift of less than 5.5 ppm. Again, consistent with amide-π hydrogen bonding, these structures show T59 exclusively involved in close contacts with aromatic rings perpendicular to the ring plane (Fig. 4g). In C36m simulations the most frequent contact involved W28 (1.8%), while W112 was the most frequently contacted sidechain by the T59 (5.7%) amide in a99sb-disp simulations (Fig. 4g).

While these amide groups are observed to interact with different parts of the protein depending on the force field (and certainly limited by sampling), we analysed the total residue-specific probability of amide-π hydrogen bonding in the native and intermediate state and found that generally tryptophan sidechains followed by tyrosine, are the most common residue types to participate in this interaction. Furthermore, these interactions occur more frequently in the intermediate compared to the native state for both force fields (Fig. 4e, Fig. S9h). Most amide-π interactions involve amides located in the β-domain and to a lesser extent the region of helices h2 and D (Fig. 4e). These regions overlap with the regions that partially unfold from their native position and can form non-native interactions with the partially exposed hydrophobic core, rendering the intermediate state rather compact and likely contributing to its stability. Together, the experiments and simulations therefore suggest that the unfolded regions sample non-native interactions with a partially exposed hydrophobic core.

## Discussion

The intermediate state of human lysozyme has long been elusive to biophysical analyses despite its direct link to hereditary systemic amyloidosis. Using an acidic condition to reduce the global cooperativity of lysozyme unfolding we previously described a pseudo-two state model revealing the exchange between the native (N) state and denatured ensemble (D)^6^. The D state comprises rapidly interconverting protein conformers with varying degrees of secondary structure denaturation. This was confirmed by the offset between unfolding curves observed via far and near-UV CD spectroscopy, alongside the absence of latent heat in calorimetric measurements. The progressive unfolding of secondary structural elements was further supported by the gradual appearance of the cross peaks in 2D HSQC spectra with increasing temperatures, but direct information on the intermediate state has remained unavailable.

In this study, we utilised CEST and CPMG NMR experiments to investigate the thermal unfolding of the I59T variant under the same experimental conditions. This enabled us to not only analyse the otherwise invisible D state at lower temperatures and monitor the entire unfolding process accurately, but most significantly, to probe and identify a distinct intermediate (I) state. The I state is in exchange with both the N and D states, is only transiently and sparsely populated (up to ∼0.6%), and expands the previous pseudo-two-state model into a modified three-state exchange model. The observed chemical shifts of the I state originate from the β-domain, C-helix and a residue from the N-terminus, which are denatured at low temperatures and exhibit highly dynamic behaviour (Fig. S5c)^7^. Such locally denatured regions are implicated in both intra- and inter-molecular interactions, initiating the aggregation of the protein into amyloid fibrils^7^.

Our all-atom unbiased MD simulations of the I state suggest that non-native hydrogen bonds and amide-π interactions account for the anomalously deshielded and shielded chemical shifts observed in the I state from CEST experiments. Specifically, the simulations indicate that the highly deshielded chemical shift of A42 arises from non-native hydrogen bonding, while I56 and T59 show a strong preference for non-native amide-π hydrogen bonding with the aromatic rings of tryptophan and tyrosine. This is qualitatively consistent with their unusually shielded chemical shifts, which deviate significantly – by ∼2.4 and ∼3.3 standard deviations, respectively – below the average chemical shifts for their residue types according to the BMRB statistics. Such deviations can be signatures of amide-ring contacts in folded proteins^27^, and have been previously observed in an SH3 folding intermediate^28^. This could indicate that amide-π hydrogen bonds are more prevalent than previously appreciated, potentially contributing to the stability of partially folded structures and folding intermediates. While classical all-atom force fields do not preclude these interactions, their stability is likely underestimated^27^. In agreement with this, we find clear evidence of such interactions in our simulations with both the C36m and a99sb-disp force fields, although the magnitudes of the predicted ensemble-averaged chemical shift changes are not entirely accurate. The limited ability of current all-atom force fields to describe these interactions, combined with insufficient sampling, may account for discrepancies between computed and experimental chemical shifts.

Our all-atom MD simulations do, however, recapitulate previously reported biophysical data on the I state, including the evidence of local unfolding in the β-domain and C-helix^6^, as well as the average folding pathway, characterised by early secondary structure formation in helices A and B and late folding of helices C and D and the β-domain^25^. The intermediate state ensemble retains stable A- and B-helices but is more molten-globule like in character due to reduced stability of tertiary interactions. While the β-domain and C-helix show the largest loss of native contacts, they remain relatively compact stabilised by non-native contacts instead of undergoing complete local unfolding.

Recent atomic structures of human lysozyme amyloid fibrils have revealed polymorphism^29,30^. Unlike the *in vitro* fibrils formed by the WT lysozyme under harsh conditions (85 ºC, 300 rpm)^29^, *ex vivo* fibrils from patients with the D87G mutation retain all four native disulfide bonds^30^. This suggests that the amyloid fibrils responsible for systemic amyloidosis are initiated by partially unfolded protein intermediates with unstructured β-domain and a native-like α-domain with intact disulfide bonds, rather than by fully denatured proteins. Our study suggests that the lysozyme intermediate is dynamic, sampling numerous non-native interactions that likely prevent complete unfolding during aggregation. The observation of amide-π hydrogen bonds, supported by the low chemical shifts of I56 and T59, highlights the role of non-native interactions in stabilising the I state and potentially contributing to the protein’s amyloidogenicity. We propose that intra-chain amide-π hydrogen bonds (spanning more than three amino acids) are more prevalent in partially folded states than in well-ordered protein structures. In folded proteins, hydrogen bonding is maximised to be energetically more favourable, and aromatic sidechains are tightly packed with hydrophobic residues in the core. In partially folded states, reduced secondary structure exposes more amides and aromatic rings, facilitating the formation of amide-π hydrogen bonds.

To prevent human lysozyme aggregation, molecules that stabilise the native state, such as ligands^30^ or nanobodies^31^, have been explored. However, given the critical role of the intermediate state in initiating aggregation into amyloid fibrils, compounds that specifically destabilise this state could also serve as effective inhibitors of amyloidosis. Aromatic compounds, for instance, might compete with residues in the protein core for amide-π interactions, thereby destabilising the I state. Our structural insights may provide a valuable foundation for designing targeted binder molecules to inhibit human lysozyme amyloid formation.

## Acknowledgements

This work is dedicated to the memory of Prof. Sir Christopher M. Dobson FRS who helped and inspired us over many years and who is missed greatly by all who knew him. We thank Ewa Klimont for assistance in protein production and purification. J.C. is supported by the Wellcome Trust (Investigator Awards 097806/Z/11/Z & 206409/Z/17/Z). J.R.K is currently supported by an MRC Career Development Award (MR/W01632X/1). J.O.S. was supported by a BBSRC London Interdisciplinary Biosciences Doctoral Programme studentship. This research was supported, in part, by the Cambridge Centre for Misfolding Diseases, and the Wellcome Trust (094425/Z/10/Z). This research was supported by the Biomolecular NMR Facility at UCL, and by the Francis Crick Institute through the provision of access to the MRC Biomedical NMR Centre. The Francis Crick Institute receives its core funding from Cancer Research UK (FC001029), the UK Medical Research Council (FC001029), and the Wellcome Trust (FC001029). We thank HWB-NMR staff at the University of Birmingham for providing open access to their Wellcome Trust-funded 900 MHz spectrometers. We acknowledge the Baskerville Tier 2 HPC service (https://www.baskerville.ac.uk/), which was funded by the EPSRC and UKRI through the World Class Labs scheme (EP/T022221/1) and the Digital Research Infrastructure programme (EP/W032244/1) and is operated by Advanced Research Computing at the University of Birmingham. This project also made use of time on HPC resources on Archer2 (ARCHER2 UK National Supercomputing service, https://www.archer2.ac.uk) granted via the UK High-End Computing Consortium for Biomolecular Simulation, HECBioSim (https://www.hecbiosim.ac.uk), supported by the EPSRC (grant no. EP/R029407/1 and EP/X035603/1). We additionally acknowledge the use of the UCL Myriad High Performance Computing Facility (Myriad@UCL), and associated support services, in the completion of this work.

