## supporting information for "Amyloid forming human lysozyme intermediates are stabilised by non-native amide-π interactions"

- Method
- Table S1
- SI movies 1-6
- Figure S1-10

### Method

All chemicals and reagents were purchased from Sigma Aldrich Ltd. (Gillingham, UK) unless otherwise stated.

**Protein expression and purification**

Lysozyme variants were expressed in *Pichia pastoris* and purified as described previously[1]. Protein purity (>95%) was confirmed using SDS-PAGE, and molecular masses were confirmed by mass spectrometry.

**Circular dichroism (CD) spectroscopy**

CD spectroscopy experiments were performed using a Jasco J-810 spectropolarimeter (JASCO Ltd, Great Dunmow, UK) equipped with a Peltier temperature controller. Protein samples (20 µM) were dissolved in phosphate buffer (50 mM, pH 1.2) and analysed in a 0.1 cm pathlength cuvette. Thermal denaturation was monitored at 222 or 270 nm with temperature increments from 5 to 95 °C (1 °C $min^{-1}$). Ellipticity values were normalized to the fraction of denatured state protein and fitted to a two-state unfolding model assuming linear baselines for both native and denatured states as described previously[2].

**NMR experiments**

*Thermal unfolding monitored by 2D NMR*

I59T human lysozyme (200 µM) was prepared in phosphate buffer (50 mM, pH 1.2) with 90% $H_2O$/10% $D_2O$. 2D $^{1}H$-$^{15}N$ HSQC spectra were recorded at temperature intervals of 2.5 or 5 °C between 5 and 60 °C on a Bruker Avance 700 MHz NMR spectrometer (University of Cambridge) and processed with NMRPipe[3] and Sparky[4]. Cross peak intensities were calculated by measuring the peak volumes using FuDA (http://www.biochem.ucl.ac.uk/hansen/fuda). The populations of the native (N) and denatured (D) states for each residue were determined by referencing the intensities of the N (at 7.5 °C) and D (57.5 °C) state, respectively[5]. Peak assignments in the $^{1}H$-$^{15}N$ HSQC spectra were performed using previously reported assignments[2].

*Peak assignments by 3D NMR*

Isotopically double-labeled ($^{13}C$, $^{15}N$) I59T was generated as described previously[2]. Backbone and side-chain resonances for the N and D states were acquired and assigned at 25 °C and 45 °C, respectively. For N state resonance assignments, CCCONH (2048 x 64 x 128 complex points and SW of 14 ($^{1}H$), 35 ($^{15}N$), 75 ($^{13}C$) ppm)[6], HCCH-TOCSY (2048 x 64 x 128, 14 ($^{1}H$), 12 ($^{1}H$), 75 ($^{13}C$) ppm)[7], HCCH-COSY (2048 x 128 x 64, 14 ($^{1}H$), 12 ($^{1}H$), 75 ($^{13}C$) ppm)[7], HNCO (2048 x 64 x 64, 14 ($^{1}H$), 30 ($^{15}N$), 12 ($^{13}C$) ppm)[8], HN(CO)CACB (2048 x 64 x 128, 14 ($^{1}H$), 30 ($^{15}N$), 62 ($^{13}C$) ppm)[8] experiments were performed on a Bruker Avance 700 MHz spectrometer (University of Cambridge). For D state resonance assignments, HNCO (2048 x 80 x 96, 15 ($^{1}H$), 22 ($^{15}N$), 10 ($^{13}C$) ppm)[8], was recorded on a 700 MHz Bruker Avance III HD (Francis Crick Institute) and H(CC)CONH (3072 x 64 x 64, 18 ($^{1}H$), 21 ($^{15}N$), 10 ($^{1}H$) ppm)[6], (H)CCCONH (3072 x 64 x 64, 18 ($^{1}H$), 21 ($^{15}N$), 10 ($^{1}H$) ppm)[6] were recorded on a 800 MHz Bruker Avance III spectrometer (University College London) equipped with a TXI cryoprobe.

*CEST NMR*

$^{15}N$ CEST experiments were performed with two $^{15}N$ $B_1$ fields (during $T_{EX}$) of 15 and 60 Hz at 11.7, 16.4, 18.8 and 22.3 T. The weak $B_1$ fields were calibrated using a sensitivity-enhanced $^{15}N$-$R_1$ measurement[9]. CEST experiments were performed at 25 - 45 °C. At 11.7 T, 42 2D $^{1}H$-$^{15}N$ HSQC spectra were acquired with a series of $^{15}N$ offsets ranging between 100.78 and 132.23 ppm obtained in increments of 0.79 ppm (40 Hz). Each 2D data set comprised 2048 x 128 complex points in the $^{1}H$ and $^{15}N$ dimensions and was recorded with 8 scans, an inter-scan delay of 1.1 s, a saturation time ($T_{EX}$) of 0.3 s. A reference spectrum was acquired with $T_{EX}$ = 0 s. At 16.4 T, 82 2D $^{1}H$-$^{15}N$ HSQC spectra were acquired with $^{15}N$ offsets ranging between 102 and 132 ppm obtained in increments of 0.38 ppm (27 Hz). Each 2D data set comprised 2048 x 128 complex points in the $^{1}H$ and $^{15}N$ dimensions and was recorded with 4 scans, an inter-scan delay of 1 s, $T_{EX}$ of 0.4 s. At 18.8 T, 62 2D $^{1}H$-$^{15}N$ HSQC spectra were acquired with $^{15}N$ offsets ranging between 100.76 and 132.36 ppm obtained in increments of 0.49 ppm (40 Hz). Each 2D data set comprised 3072 x 128 complex points in the $^{1}H$ and $^{15}N$ dimensions and was recorded with 4 scans, an inter-scan delay of 1.3 s, $T_{EX}$ of 0.35 s. At 22.3 T, 82 2D $^{1}H$-$^{15}N$ HSQC spectra were acquired with $^{15}N$ offsets ranging between 102 and 132 ppm obtained in increments of 0.38 ppm (37 Hz). Each 2D data set comprised 2048 x 128 complex points in the $^{1}H$ and $^{15}N$ dimensions and was recorded with 8 scans, an inter-scan delay of 0.95 s, $T_{EX}$ of 0.4 s.

$^{13}C$ CEST experiments for methyl carbons were performed at 35 °C, 22.3 T, using a $^{13}C$ $B_1$ field strength of 25 Hz with $T_{EX}$ = 0.3 s. 81 2D $^{1}H$-$^{13}C$ HSQC spectra were acquired with $^{13}C$ offsets ranging between 12.23 and 31.05 ppm obtained in increments of 0.21 ppm (50 Hz). Each 2D data set comprised 4096 x 220 complex points in the $^{1}H$ and $^{13}C$ dimensions and was recorded with 4 scans, an inter-scan delay of 1.6 s, $T_{EX}$ of 0.3 s. $^{13}C_{\alpha}$ CEST experiments were performed at 35 °C, 18.8 T, using a $^{13}C$ $B_1$ field strength of 30 Hz with $T_{EX}$ = 0.3 s. 77 2D $^{1}H$-$^{13}C$ HSQC spectra were acquired with $^{13}C$ offsets ranging between 50.43 and 69.06 ppm obtained in increments of 0.25 ppm (50 Hz). Each 2D data set comprised 3072 x 256 complex points in the $^{1}H$ and $^{13}C$ dimensions and was recorded with 4 scans, an inter-scan delay of 1 s, $T_{EX}$ of 0.3 s.

$^{1}H$ CEST experiments for methyl sidechains were performed at 35 °C, 22.3 T, using a Z-Z exchange-based pulse sequence with a $^{1}H$ $B_1$ field strength of 60 Hz with $T_{EX}$ = 0.3 s. 145 2D $^{1}H$-$^{13}C$ HSQC spectra with $^{1}H$ offsets ranging between -2.84 and 4.73 ppm obtained in increments of 0.05 ppm (50 Hz). Each 2D data set comprised 4096 x 200 complex points in the $^{1}H$ and $^{13}C$ dimensions and was recorded with 4 scans, an inter-scan delay of 1s. $^{1}H_{\alpha}$ CEST experiments were performed at 35 °C, 22.3 T with a $^{1}H$ $B_1$ field strength of 45 Hz with $T_{EX}$ = 0.3 s. 82 2D $^{1}H$-$^{13}C$ HSQC spectra with $^{1}H$ offsets ranging between 2.96 and 5.94 ppm obtained in increments of 0.04 ppm (35 Hz). Each 2D data set comprised 4096 x 96 complex points in the $^{1}H$ and $^{13}C$ dimensions and was recorded with 4 scans, an inter-scan delay of 1 s.

*CPMG relaxation dispersion NMR*

CPMG relaxation dispersion profiles were recorded at 16.4, 18.8 and 21.1 T at 37.5 and 45 °C as described previously[10]. A constant-time CPMG interval (40 ms) with $^{1}H$ continuous-wave decoupling along with 20 $\nu_{CPMG}$ values ranging from 25 to 1000 Hz were used. Each 2D data set comprised 2048 x 128 complex points in the $^{1}H$ and $^{15}N$ dimensions and was recorded with 8 scans, an inter-scan delay of 1 s.

*Rotating frame ($R_{1\rho}$) relaxation dispersion NMR*

On- and off-resonance $R_{1\rho}$ measurements were performed at 37.5 °C for the denatured peak of G129 at a field strength of 16.4 T as previously described[11]. $^{15}N$ RF field strengths were calibrated by pulse nutation experiments. On-resonance $R_{1\rho}$ experiments were performed at spin-lock field strengths ($\omega_1/2\pi$) ranging from 100 to 2000 Hz. Off-resonance experiments were recorded at three spin-lock field strengths (200, 1000 and 2000 Hz) with 12 to 20 offsets. Each data set comprised 2048 complex points in the $^{1}H$ dimension and was recorded with 64 scans, an inter-scan delay of 1 s.

**NMR data fitting**

$^{15}N$ CEST and CPMG data were processed using FuDA and fitted using the ChemEx software package (http://www.github.com/gbouvignies/chemex). For two-state model, CEST data from 109 residues in both the N and D states were globally fitted to a two-state unfolding model (N-D) at 25 – 37.5 °C (Fig.1e). For three-state model, both the N and D state CEST and CPMG data from 15 residues (Fig.S2) were globally fitted to a triangular model as well as three sequential models (N-I-D, N-D-I, I-N-D). On- and off-resonance $R_{1\rho}$ data were globally fit to the Trott and Palmer equation[12]:

$$R_{1\rho} = R_1 cos^2\theta + (R_2 + R_{ex}) sin^2\theta$$

where

$$R_{ex} = \frac{p_B \Delta\omega^2 k_{ex}}{(\delta_A + \Delta\omega)^2 + {\omega_1}^2 + {k_{ex}}^2}$$

with $\omega_1$ is the 15N spin-lock field strength, $\delta_A$ is the $^{15}N$ resonance offset of the A state and $\Delta\omega$ is the frequency difference between state A and B in the asymmetric population limit ($p_A \gg p_B$).

**MD simulations methods**

*Ratchet-and-pawl MD (rMD) simulations*

All simulations in this study were run using GROMACS (v2018 and v2021)[13]. Starting structures for rMD folding simulations were generated using short, all-atom structure-based model[14] simulations where all native contacts were deleted, resulting in rapid unfolding and the generation of random sterically allowed conformers. All four disulphide bridges were kept oxidised. Resulting structures were centred in a dodecahedron box with a padding of 1.0nm. Following solvation and neutralisation using chlorine ions, structures were energy minimised using the steepest-descent algorithm and equilibrated for 500ps first in the NVT (308K) and then NPT (308K, 1bar) ensemble using heavy atom position restraints with a force constant of 1000 kJ $mol^{-1}$ $nm^{-1}$. Subsequently, the structures were relaxed without position restraints in the NPT ensemble (308K, 1bar) for 1ns. The LINCS algorithm [15] was used to constrain all bonds to hydrogens together with a timestep of 2fs and leap-frog algorithm for integration. The temperature was controlled using the velocity rescaling algorithm [16] with a time constant of 0.1ps. During equilibration, the pressure was maintained using the Berendsen barostat [17] with a compressibility of $4.5x10^{-5}$ $bar^{-1}$ and the Parrinello-Rahman algorithm during the 1ns relaxation and following production rMD simulations [18]. Simulations were parameterised using the CHARMM36m (C36m) [19] and a99sb-disp [20] force fields. To mimic the low pH conditions of the NMR experiments, all titratable groups were protonated accordingly. Nonbonded van der Waals interactions were treated to a cut-off at 1.2nm (with a switching function from 1.0nm for C36m). Electrostatics were calculated to a cut-off of 1.2nm and long-range electrostatics were computed using the Particle Mesh Ewald (PME) method [21].

Production simulations employing the rMD algorithm were run for 5ns with a 2fs timestep at 350K and 1bar in the NPT ensemble. In rMD simulations a history-dependent biasing force is introduced when the protein attempts to backtrack along its folding trajectory defined as [22,23]

$$F_{rMD} = \begin{cases} \frac{\kappa}{2}\left(\rho(t) - \rho_m(t)\right)^2, \rho(t) > \rho_m(t) \\ 0, \rho(t) \le \rho_m(t) \end{cases}$$

where $\rho(t) = \left(CV(t) - CV_{target}\right)^2$ and $\rho_m(t) = \min_{0\to t}(CV)$ . $\kappa$ is the force constant controlling the strength of the biasing force applied, $CV(t)$ is the value of a chosen collective variable at time t and $CV_{target}$ is the defined target value of the collective variable (in this case the native state). We used a $\kappa$ value of $1x10^{-9}$ kJ $mol^{-1}$. The collective variable (CV) for ratcheting used here is the difference in the contact map of the structure at time t, $C_{ij}(X)$, and the native state, $C_{ij}(X_{native})$, defined as [24,25]

$$CV = \sum_{|i-j|>35} \left(C_{ij}(X) - C_{ij}(X_{native})\right)^2$$

$$C_{ij}(X) = \frac{1 - \left(\frac{r_{ij}}{r_0}\right)^6}{1 - \left(\frac{r_{ij}}{r_0}\right)^{10}}$$

All heavy atom indices i and j are included in the contact map and $r_{ij}$ is the distance between heavy atoms while $r_0$ is 0.75nm. All distances up to and including 1.2nm in the native crystal structure (PDB 2MEH[26] ) are included. We simulated a total of 36 folding trajectories (each 5ns in length) with different random initial velocities for 50 different input structures with each force field, yielding a total of 9μs of rMD data per force field.

To calculate the kinetic folding free energy landscape, we extracted an ensemble of folding trajectories that reached the native state (defined as Cα-RMSD≤3.0Å relative to the crystal structure). For these ensembles of folding trajectories, we estimated the free energy landscape as a function of Cα-RMSD and Q (the fraction of native contacts, defined as in [27] with λ=1.5), smoothed by a gaussian filter with σ=2. The intermediate basin was defined as a local minimum with structures below 11 kJ/mol. We then clustered all structures from the intermediate basin using the GROMOS algorithm [28], the Cα-RMSD as a similarity measure and 5Å as cut-off value using.

*Unbiased MD simulations of the intermediate state*

We chose two starting structures per force field from the intermediate clusters where the C-helix is unfolded and lacks tertiary contacts with the β-domain. Each starting structure was centred in a dodecahedron box with 1.2nm of padding and solvated/neutralised as before for rMD simulations. We parameterised the simulations with the respective force fields and protonated all titratable groups. Simulation boxes were energy minimised by the steepest descent algorithm and then equilibrated for 500ps in the NVT ensemble (308K) followed by 500ps in the NPT ensemble (308K, 1bar). We employed the same simulation algorithms and cut-off values as for the rMD simulations. To facilitate sampling of longer timescales, we ran simulations using hydrogen mass repartitioning (HMR) [29], allowing a timestep of 4fs. Four production simulations of 10μs were launched for each starting structure with different initial velocities. This yielded a total of 160μs of MD for the intermediate state. Control simulations for each force field of the native state were initiated from the crystal structure[26] and run equivalently for 4x2.5μs with different initial velocities.

*Ensemble analysis*
We used the python packages MDAnalysis [30] and MDTraj [31] for analysis of the ensembles involving atomic coordinates. The fraction of native contacts was calculated as defined by Best *et al.* [27]

$$Q(X) = \frac{1}{N}\sum_{ij}\frac{1}{1+e^{\left(\beta\left(r_{i,j}-\lambda r^{0}{}_{i,j}\right)\right)}}$$

where $r_{i,j}$ and $r^{0}{}_{i,j}$ are the distances between atoms $i$ and $j$ in frame $X$ and the template structure, respectively, $\beta$ modulates the smoothness of the switching function (default value $5\text{Å}^{-1}$used) and $\lambda$ is a factor allowing for fluctuations of the contact distance (set to 1.5).

Contact maps were calculated using distances between all heavy atoms and a cut-off distance of 5.0Å. Secondary structure populations were computed with the DSSP algorithm [32] with the implementation in MDTraj. GROMACS [13] was used to calculate the solvent-accessible surface area (SASA) using all protein atoms. Backbone chemical shifts were calculated from MD ensembles using SHIFTX2, which has the highest reported accuracy [33]. To compare the global agreement with chemical shifts from different nuclei, we calculate a chemical shift score defined as the summed RMSD for each nuclei normalised by the average error of the method for the respective nucleus. Finally, amide-π hydrogen bonds were defined by the following criteria:

1. An amide proton to ring center of mass (COM) distance of less than 4.5Å
2. The angle between the ring normal vector and vector formed by the amide proton and ring COM must be less than 54.7 degrees (i.e., the amide is perpendicular to the ring plane)
3. The angle formed by the amide nitrogen, proton and ring COM must be higher than 120.0 degrees (hydrogen bonding directionality)
4. The amide proton is not involved in another non-aromatic hydrogen bond (normal hydrogen bonds are defined by amide proton – acceptor distances of less than 2.5Å and angles higher than 120.0 degrees).

**MD simulations supplementary text**
*Folding process*
To validate the average folding pathway observed in the rMD simulations, we compared our folding simulations to biophysical data available in the literature. First, we assessed the order of secondary structure formation, focusing on the main helices A, B, C, D, and the β-sheet

(**Figure S8c-d**). Pulse deuterium labelling studies revealed that during non-oxidative refolding of HuL helices A and B are protected rapidly from exchange, before the other secondary structure elements [34]. This is followed by formation of secondary structure in helices C and D, and afterwards in the β-sheet [34]. Consistent with these experiments, we observe for both force fields that the β-sheet forms the lowest amount of secondary structure on average in our folding simulations, and it forms more slowly than the main helices (**Figure S8d**). Furthermore, helix D is predicted to form less structure and more slowly than helices A and B, also consistent with experiments. However, helices A-C appear to form similar amounts of structure and follow similar kinetics on average, suggesting in simulations helix C is too stable and forms too early. We also assessed the order of tertiary contact formation of the main helices and β-sheet, which showed that for both force fields the simulations predict rapid collapse of formation of tertiary native contacts for all structural elements, except helix C which is delayed (**Figure S8e**). This observation is consistent with previous NMR data showing that the intermediate observed during thermal unfolding mutant has an unfolded C-helix and β-sheet [35].

*Trp residues SASA*

To assess the accuracy of tertiary structure formation in our simulations further, we computed the solvent accessibility of two tryptophan indole groups (Trp109 and Trp112), which were seen by pulse labelling NMR to follow different protection kinetics. Consistent with experiments [34], the simulations predict more rapid burial of Trp112 than Trp109 (**Figure S8f-g**). This is also consistent with late folding of the C-helix against the rest of the protein, since Trp112 reports mainly on contacts between helices D and B and Trp109 on contacts with the C-helix (**Figure S8f**). Lastly, stopped-flow CD experiments showed that during refolding ~50% of native like signal in the near UV range develops rapidly, followed by slower development of the remaining native signal [34]. This suggests that two out of the four buried tryptophan residues are buried prior to burial of the other two. Indeed, this behaviour is also predicted by our simulations with both force fields, showing that Trp28 and Trp109 are buried more slowly than Trp64 and Trp112 (**Figure S8h**). This is consistent with Trp28 and Trp109 reporting on tertiary contacts with the C-helix and partial exposure of these sidechains in the intermediate state.

*Secondary structure stability*

Next, we assessed the stability of secondary structure in the unbiased MD simulations and found that helices A and B maintained the most helical structure, while helix C was more variable and can exhibit large amounts of helicity but also almost complete unfolding. Helix D also has a reduced helicity (**Figure 4d,S9c**). This is broadly consistent with the C-helix forming at least some secondary structure before the β-domain during refolding [34], and thus before tertiary contacts are established, but also partial unfolding of helix C in the intermediate state. The β-domain, in contrast to the native state, shows a significant reduction in the amount of β-sheet structure, consistent with unfolding of this domain (**Figure 4d,S9c**).

**Table S1**. $C_\alpha$ and $C_\beta$ chemical shifts of cysteine residues in the native and denatured states at 35 ºC. All native cysteine residues are oxidised as expected. $C_\beta$ chemical shifts of C6, C65 and C95 confirm that three disulfide bonds (C6-C128, C65-C81, C77-C95) are intact (oxidised)[36,37]. The chemical shifts for C30 and C116 could not be assigned (n/a) due to overlapping peaks in the denatured state spectra. However, their disulfide bond is predicted to be intact, as the C77-C95 bond (the weakest of the four[38]) remains intact.

| residue | $C_\beta$ | | $C_\alpha$ | |
|---|---|---|---|---|
| | native | denatured | native | denatured |
| 6 | 32.861 | 40.378 | 54.19 | 56.388 |
| 30 | 44.873 | n/a | 60.646 | n/a |
| 65 | 45.481 | 41.272 | 53.087 | 56.37 |
| 77 | 39.277 | n/a | 55.933 | n/a |
| 81 | 37.715 | n/a | 56.846 | 56.93 |
| 95 | 34.979 | 41.924 | 55.3 | 56.551 |
| 116 | 44.066 | n/a | 54.664 | n/a |
| 128 | 34.931 | n/a | 51.92 | 55.632 |

**SI Movies** 1-6.
S1. N state a99sb-disp
S2. N state C36m
S3. I state a99sb-disp (1 input structure, 4x10us)
S4. I state a99sb-disp (1 input structure, 4x10us)
S5. I state C36m (1 input structure, 4x10us)
S6. I state C36m (1 input structure, 4x10us)

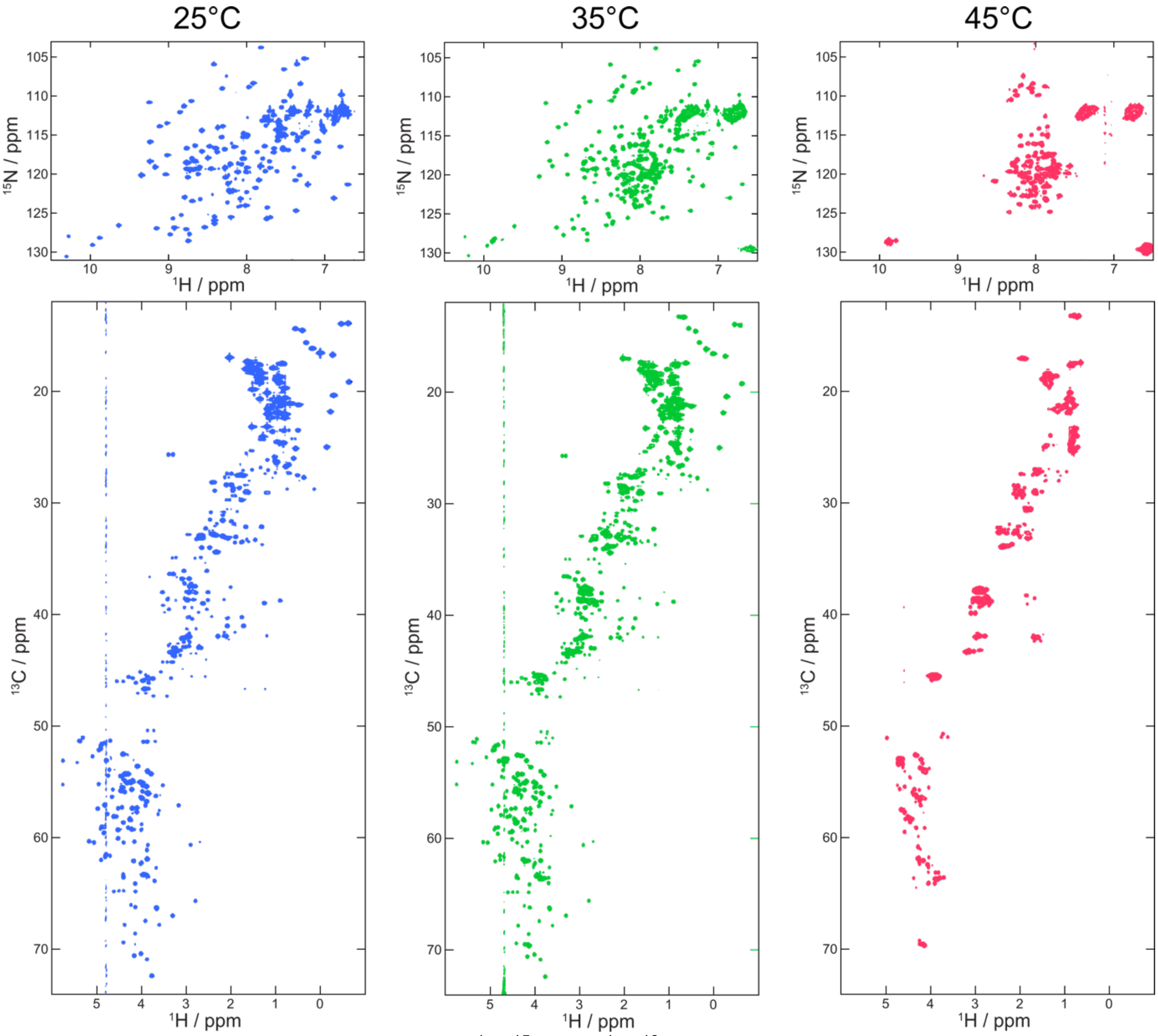


**Figure S1.** Thermal unfolding of I59T. 2D $^{1}H$-$^{15}N$ and $^{1}H$-$^{13}C$ spectra of I59T at pH1.2 were recorded at 25, 35, 45 ºC on an 800 MHz spectrometer.

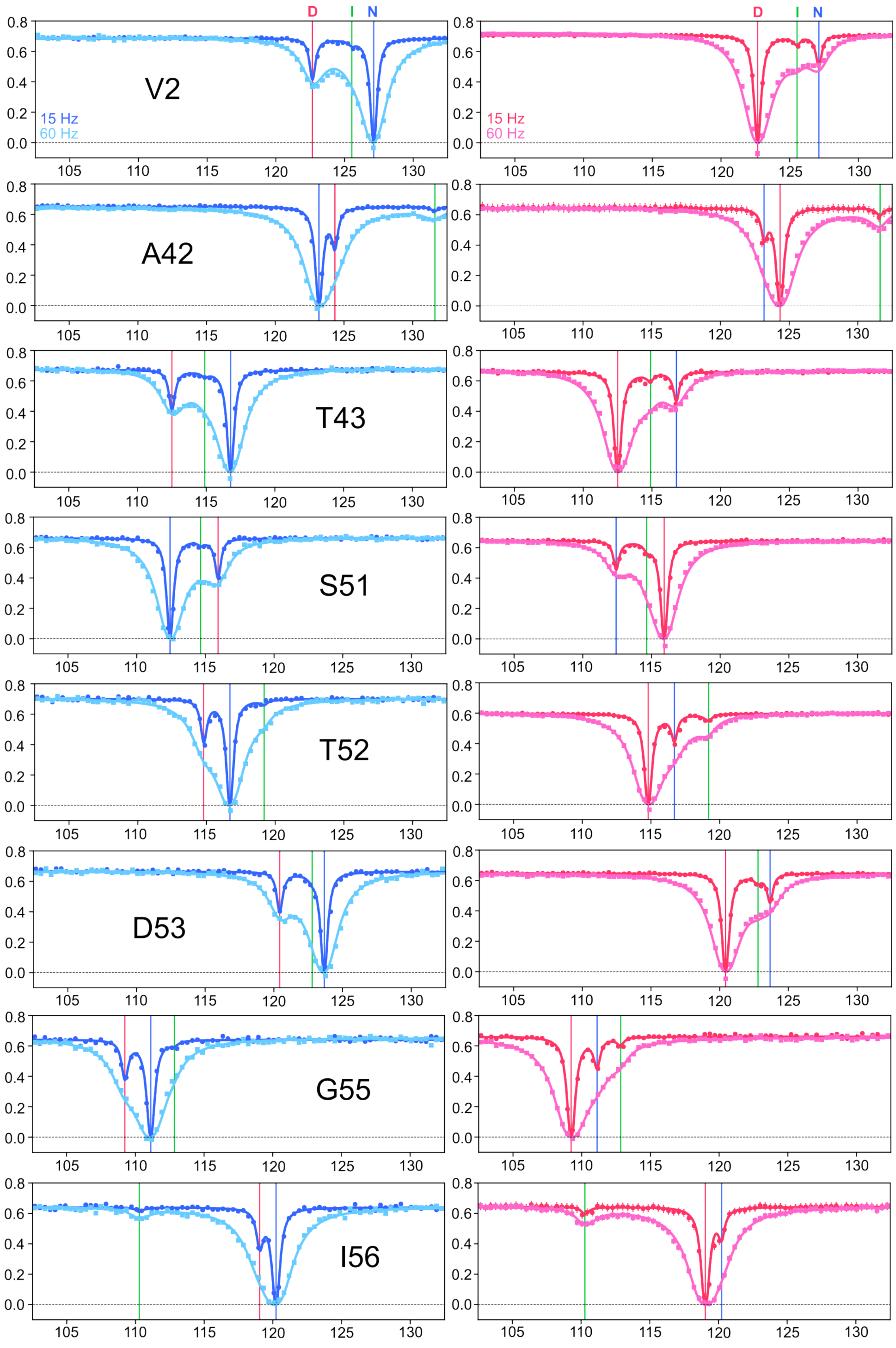
D
I
N
V2
A42
T43
S51
T52
D53
G55
I56
15 Hz
60 Hz

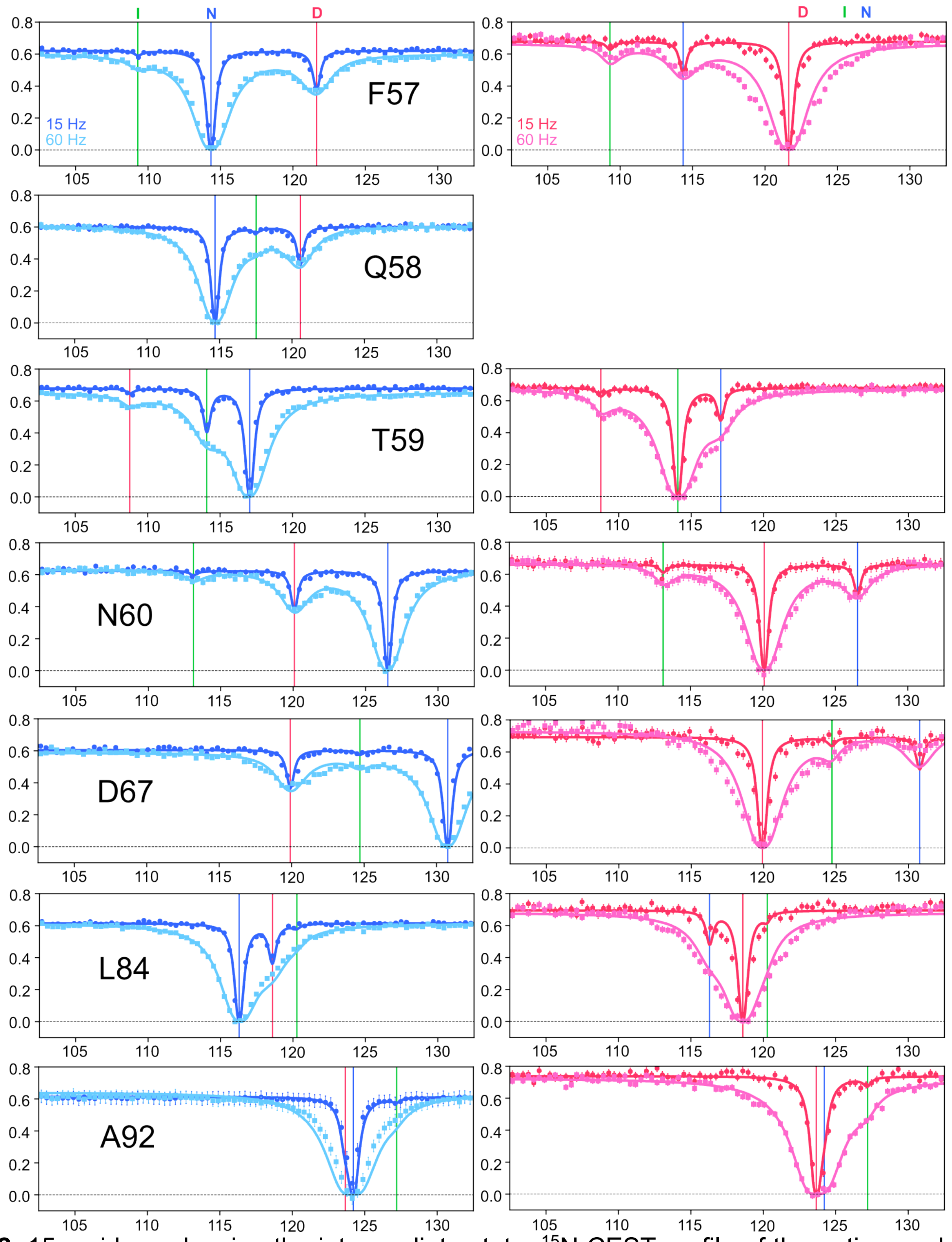


**Figure S2.** 15 residues showing the intermediate state. $^{15}$N CEST profile of the native peaks at 15Hz B1 field recorded at 800 MHz at 35 ºC. Blue, red and green vertical lines indicate the chemical shift of the native, denatured and intermediate states, respectively. The denatured peak of Q58 is not included due to a significant overlap with other peaks.

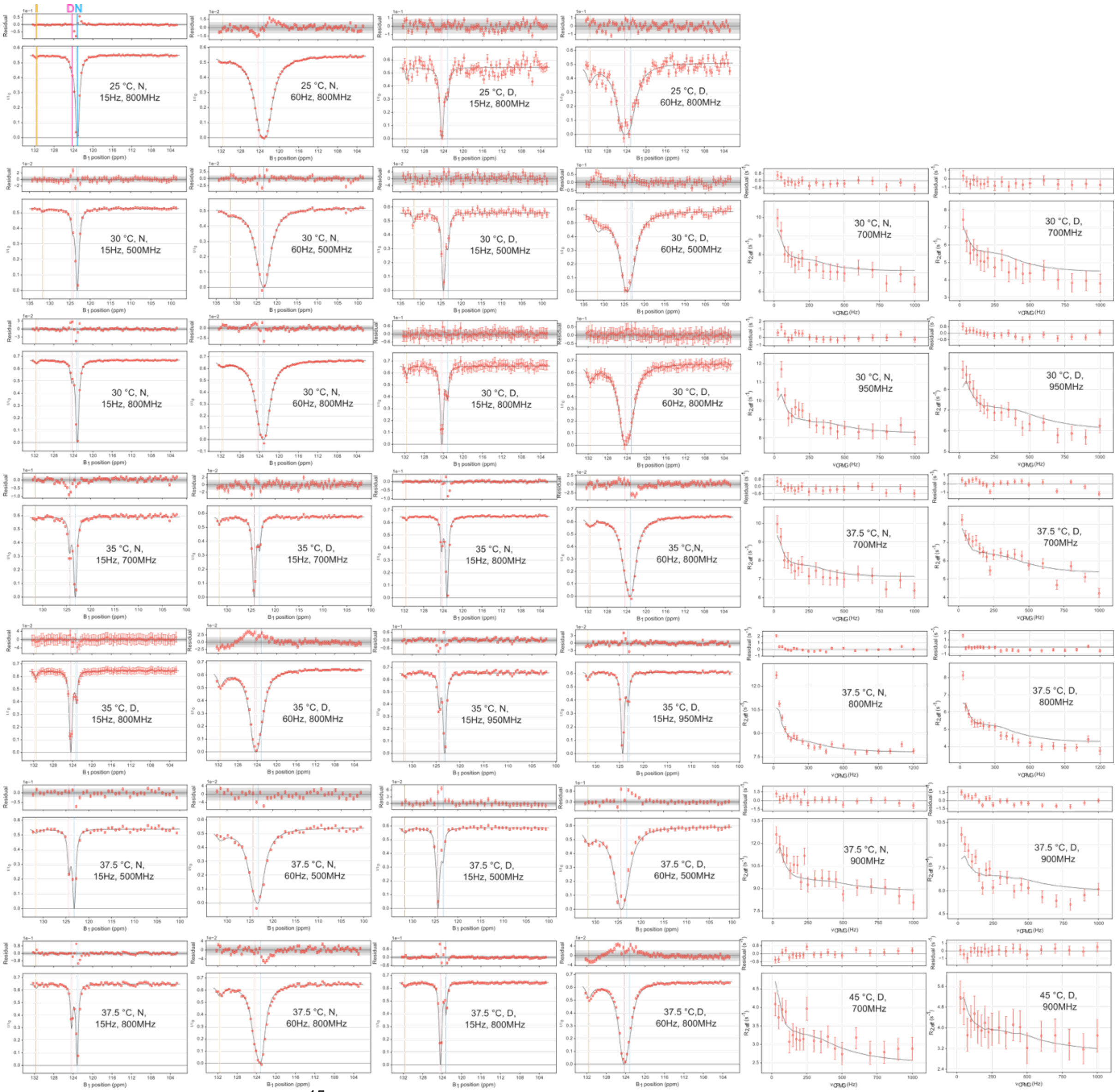


**Figure S3.** Global fitting of $^{15}$N CEST and CPMG RD data to a three-state unfolding model. Data for residue A42 collected at various temperatures and field strengths used for ChemEx data fitting are shown as an example. N and D represent the data from the native and denatured peaks. Blue, red and yellow solid vertical lines in CEST profiles represent the chemical shifts of the native, denatured and intermediate states, respectively.

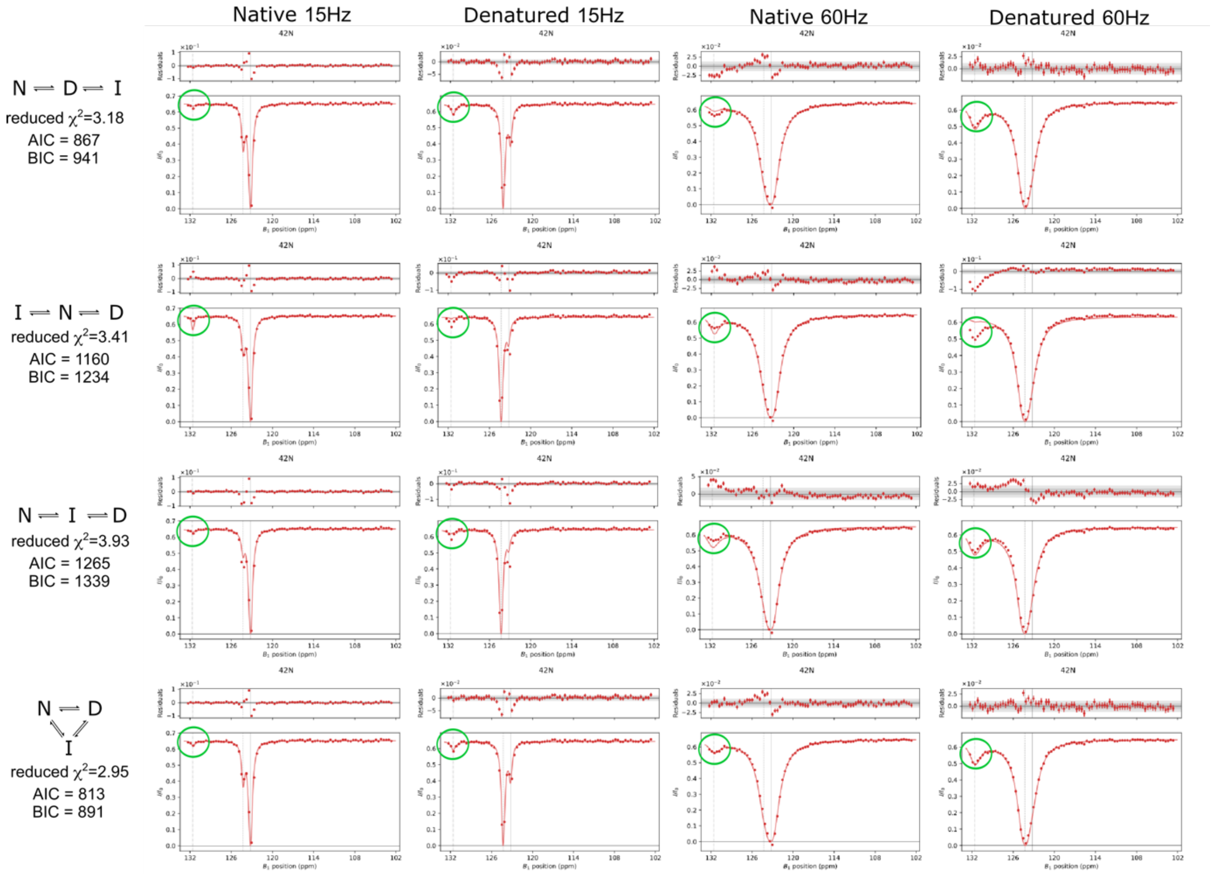


**Figure S4.** Global fitting of $^{15}N$ CEST and CPMG data of A42 to four three-state exchange models. The native and denatured peak CEST profiles recorded with 15 and 60 Hz B1 field at 800 MHz and 35 ºC are shown for comparison. Green circles indicate the CEST dips corresponding to the intermediate (I) state. The triangular model (bottom) shows the best fit, as evidenced by the lowest $\chi^2$, AIC and BIC values.

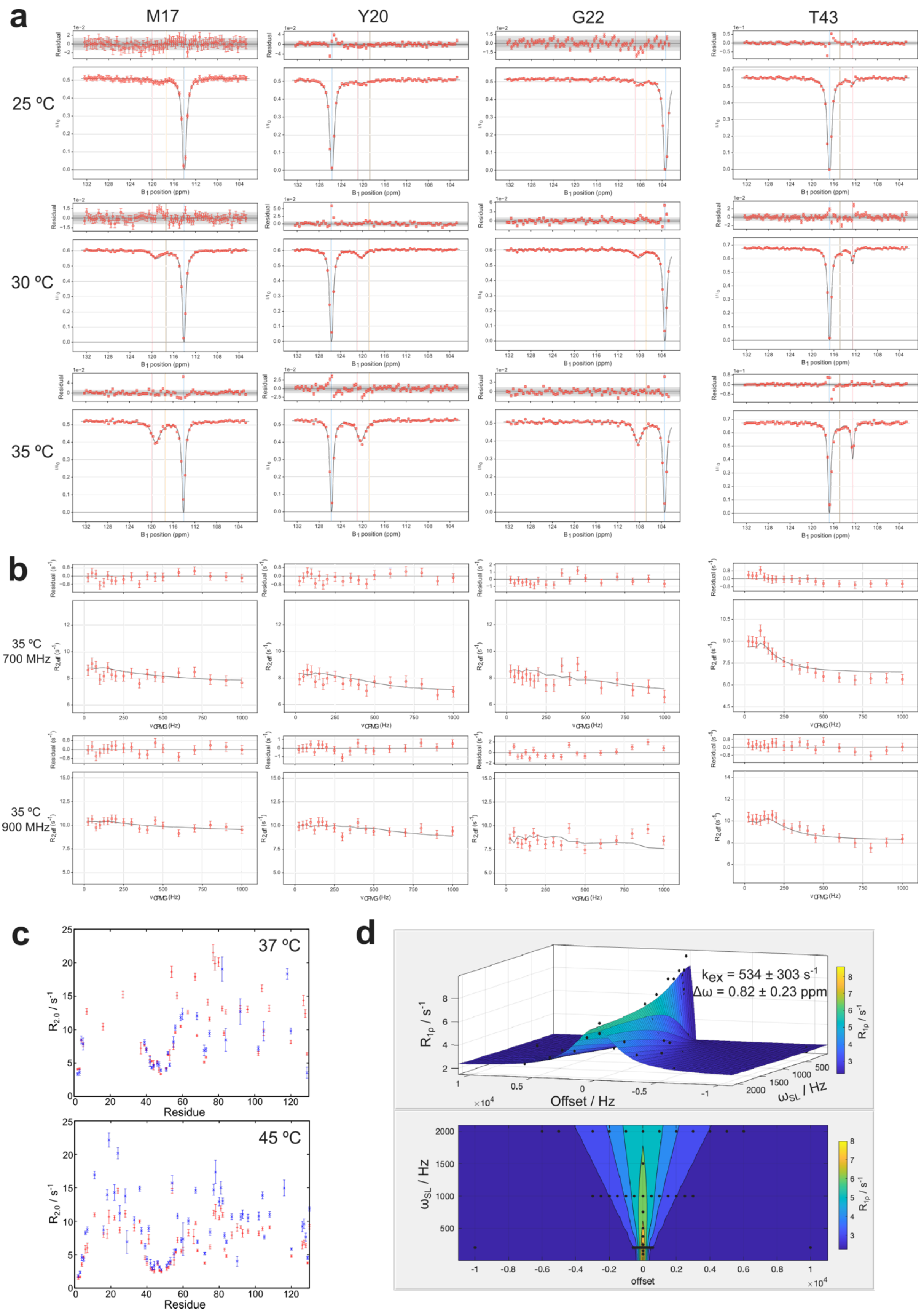


**Figure S5.** Probing the dynamics of α-domain residues. CEST (a) and CPMG (b) data the N state peaks of α-domain residues (helix A) fitted to a three-state unfolding model. Three residues (M17, Y20 and G22) from helix A show distinct behaviour compared to the β-domain residue (T43). CEST data are recorded at 800 MHz. (c) $R_{2,0}$ values from the fitting of CPMG data for D state peaks at 37 (top) and

45 (bottom) ºC, recorded at 700 (red) and 900 (blue) MHz. (d) On- and off-resonance $R_{1\rho}$ relaxation dispersion data for the denatured peak of G129 were fit to the Trott and Palmer equation[12]. The exchange rate ($k_{ex}$) was 534 $\pm$ 303 $s^{-1}$. Experiments were recorded at 37.5 ºC at 700 MHz.

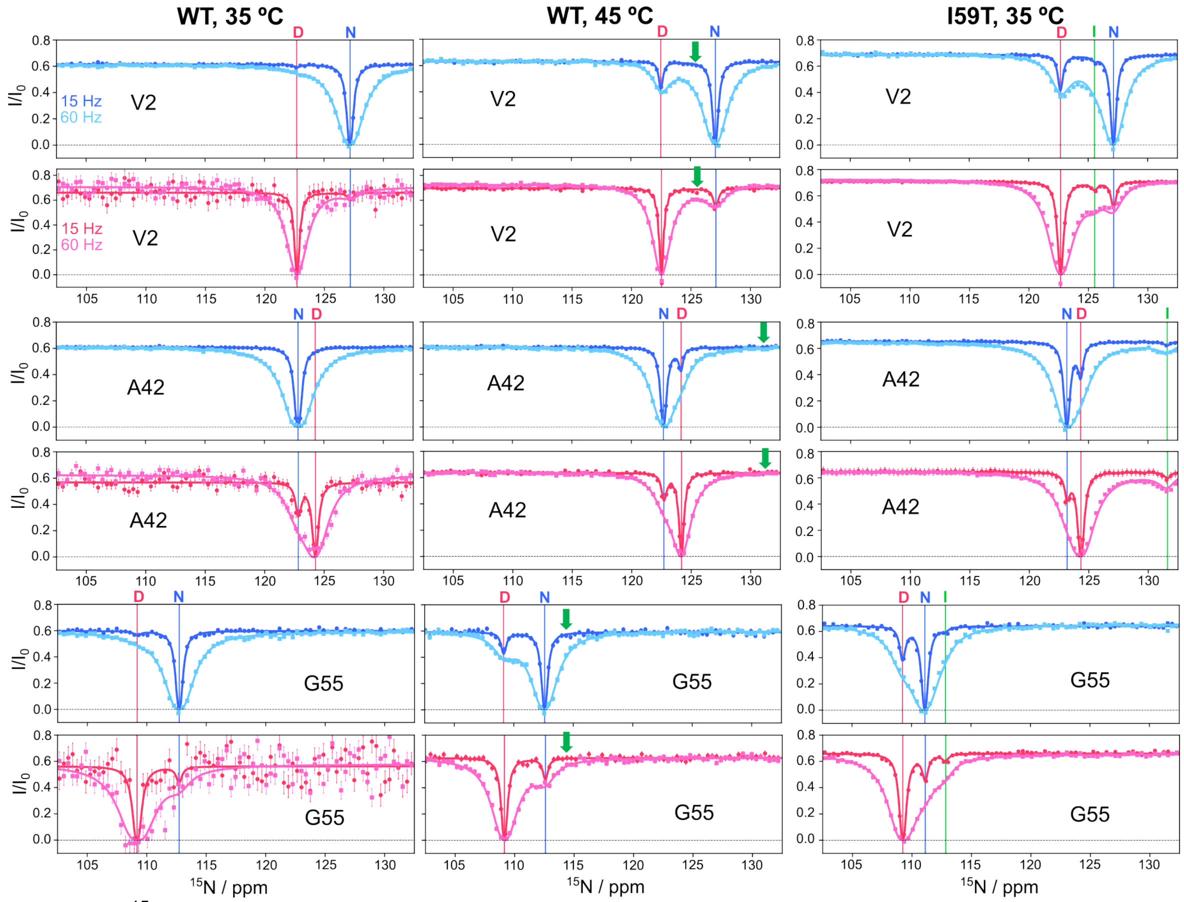


**Figure S6.** $^{15}$N CEST data for V2, A42 and G55 in the WT (35 and 45 ºC) and I59T (45 ºC). Red and Blue data points show the CEST data from the native and denatured peaks, respectively. Green arrows highlight the absence of intermediate state CEST dips in the WT data. All spectra were recorded at 800 MHz. Blue, red and green vertical lines indicate the chemical shift of the native, denatured and intermediate states, respectively.

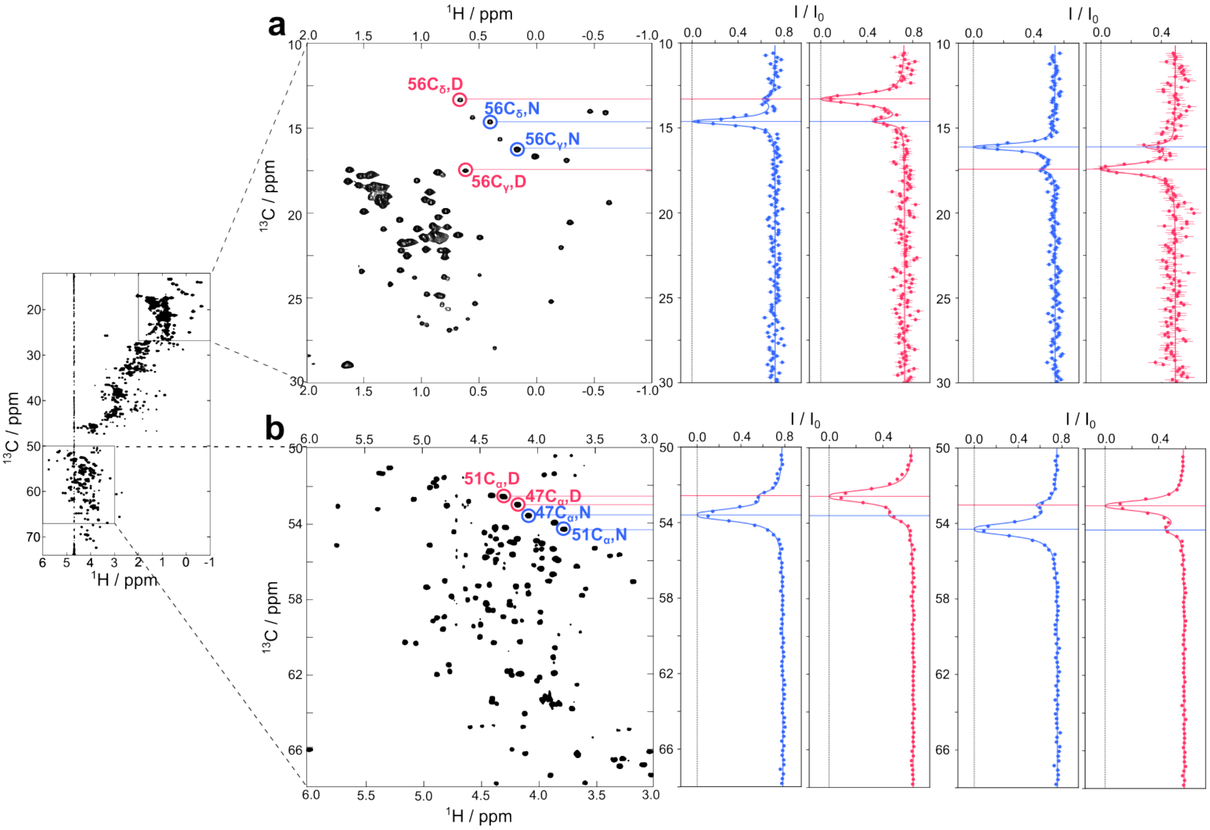


**Figure S7.** $^{13}$C CEST of I59T at 35 ºC. (a) $^{13}$C CEST on 56C$_\gamma$ and 56C$_\delta$ recorded at 950 MHz. (b) $^{13}$C CEST on 47C$_\alpha$ and 51C$_\alpha$.

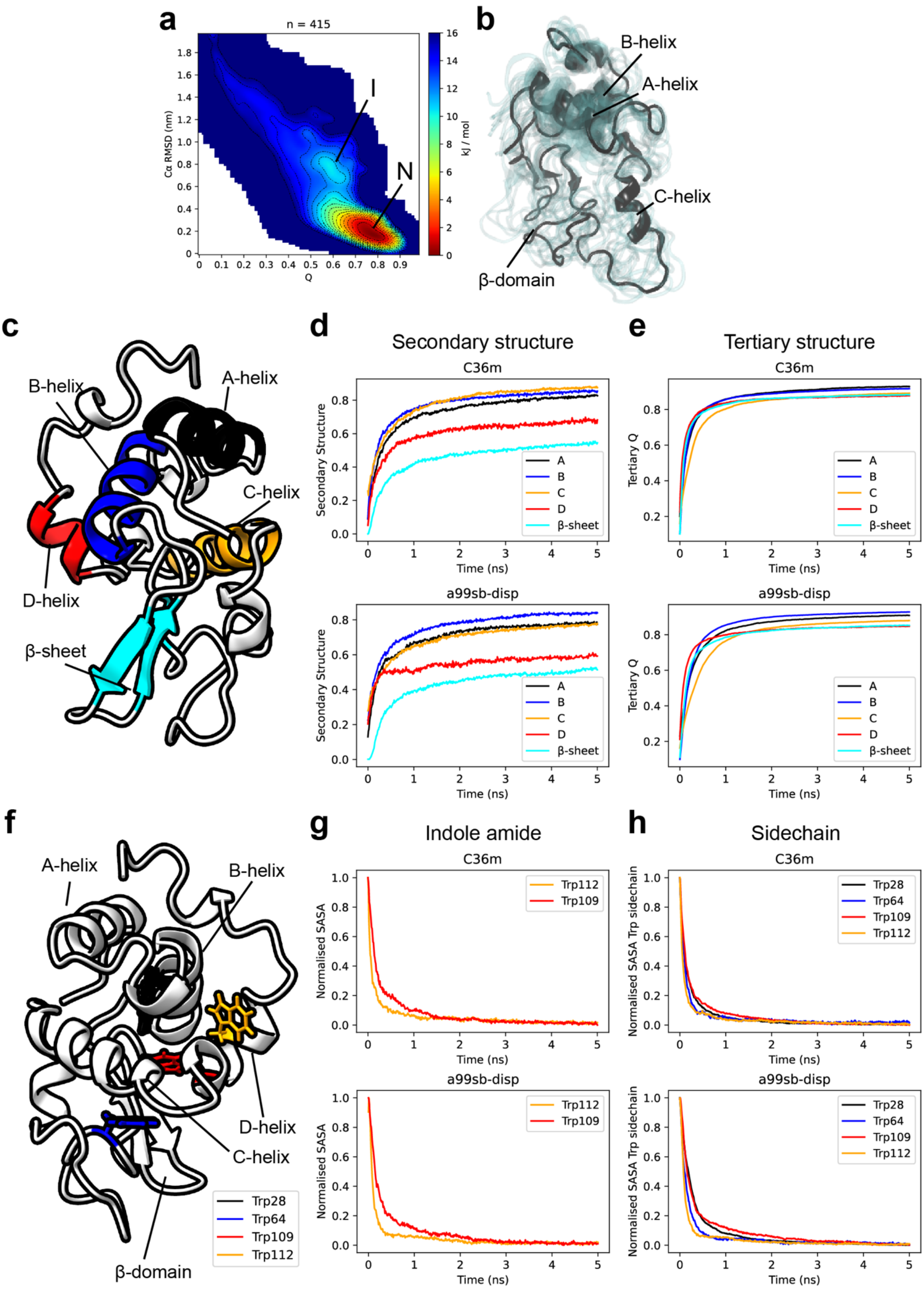


**Figure S8.** Analysis of the folding free energy landscape and pathway observed in rMD simulations. (a) Kinetic folding free energy landscape of HuL I59T calculated from all (n = 415) rMD trajectories that reach the native state with the C36m force field. The regions of the native state and metastable intermediate are annotated. (b) Representative ensemble of the folding intermediate with an unfolded C-helix and β-domain from panel a. With C36m, this state is only the second most populated cluster, in contrast to a99sb-disp (the most populated cluster obtained with C36m corresponds to an unfolded helix A, which is not supported by experiments). (c) Crystal structure (PDB 2MEH)[26] annotated with the main secondary structure elements. (d) Average fraction of secondary structure formed in each secondary structure element as a function of time for all rMD trajectories that reached the native state. Secondary structure populations were calculated using DSSP (see methods). (e) Average fraction tertiary contacts, Q (native contacts of each secondary structure element with the rest of the protein, see methods), as a function of time for all rMD trajectories that reached the native state. (f) Crystal structure annotated with the four buried tryptophan residues. (g) Average time course of the tryptophan indole (atoms NE1 and HE1) solvent-accessibility group for the indicated residues from all rMD trajectories that reached the native state. The SASA was normalised by the minimum and maximum values observed to account for intrinsic differences between the two tryptophan residues in the native state. (h) SASA time courses for all four buried tryptophan sidechains.

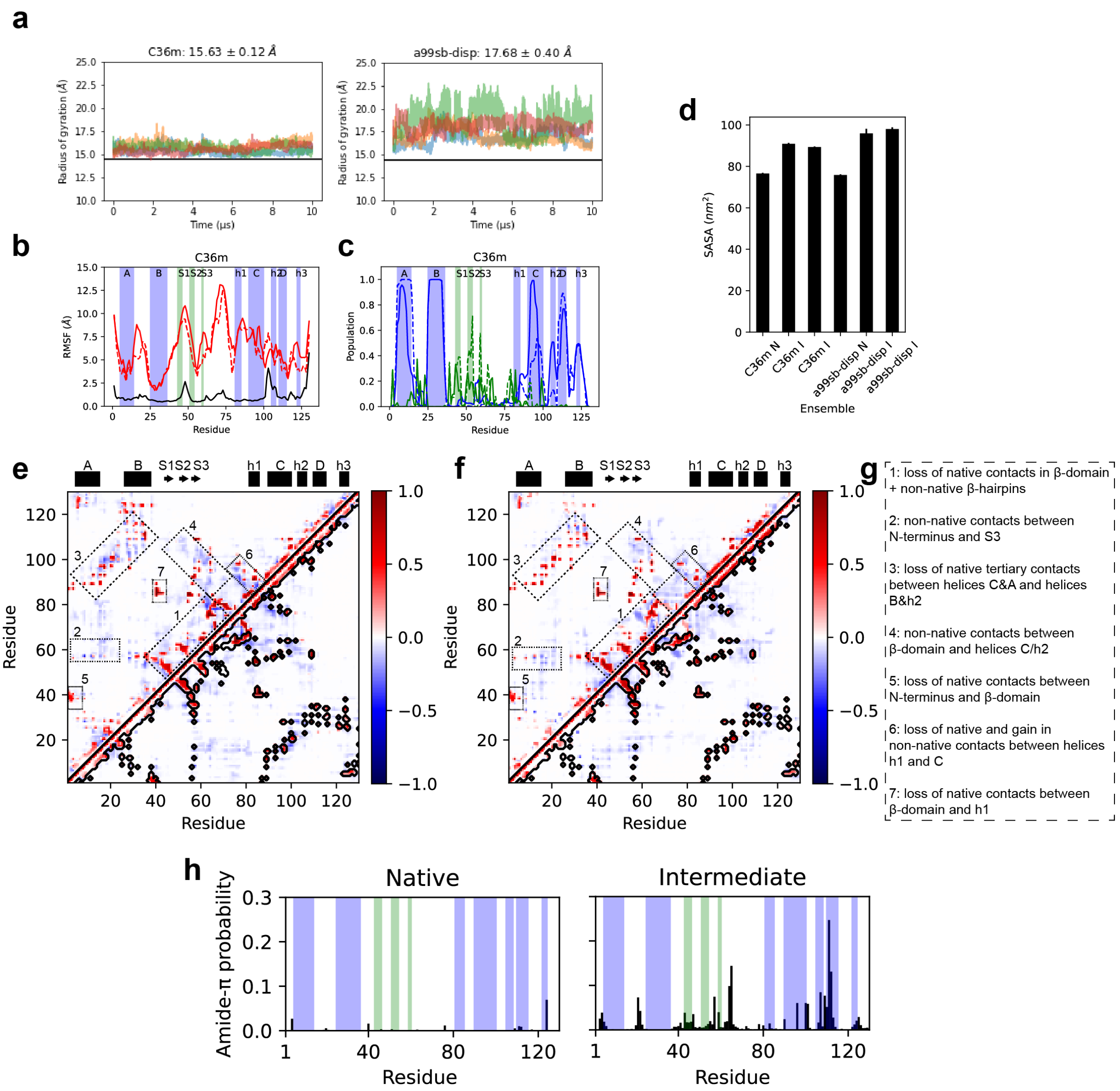


**Figure S9.** Conformational dynamics of the intermediate state ensemble sampled by unbiased MD simulations. (a) Stability of intermediate states in unbiased, long-timescale MD simulations assessed by fluctuations in the radius of gyration for both force fields. Each plot contains 8x10μs trajectories generated from two starting structures (four simulations each). The horizontal line represents the radius of gyration obtained from the native state MD simulations (average ± standard error from 4x2.5μs). (b) Backbone dynamics of the native (black) and intermediates state (red) calculated from the unbiased MD simulations for the C36m force field, quantified by the RMSF (Cα atoms). The solid and dashed red lines represent the RMSF calculated from 4x10μs of unbiased MD simulations for each intermediate starting structure to assess the reproducibility of the structural ensemble. (c) Average secondary structure populations (α-helix – blue, β-sheet – green) of the intermediate state ensemble obtained with the C36m force field (solid and dashed line represent the 4x10μs ensembles from two starting structures, respectively). (d) Average solvent-accessible surface area (SASA) of the native state (N, 4x2.5μs) and intermediate state ensembles for both force fields (I, 4x10μs). (e) Difference contact map of the intermediate state (native – intermediate) for the intermediate state ensemble obtained with the C36m force field. Contacts were defined when heavy atoms of two residues come within 5.0Å of each other. Red represents the loss of native contacts and blue the gain in non-native contacts. The black contour is the contact map of the native state ensemble. (f) Difference contact map of the intermediate state obtained with the a99sb-disp force field. (g) Description of highlighted regions in the contact maps in panels e-f. (h) Ensemble-averaged probability of forming amide-π interactions in the native and intermediate state ensemble obtained with C36m.

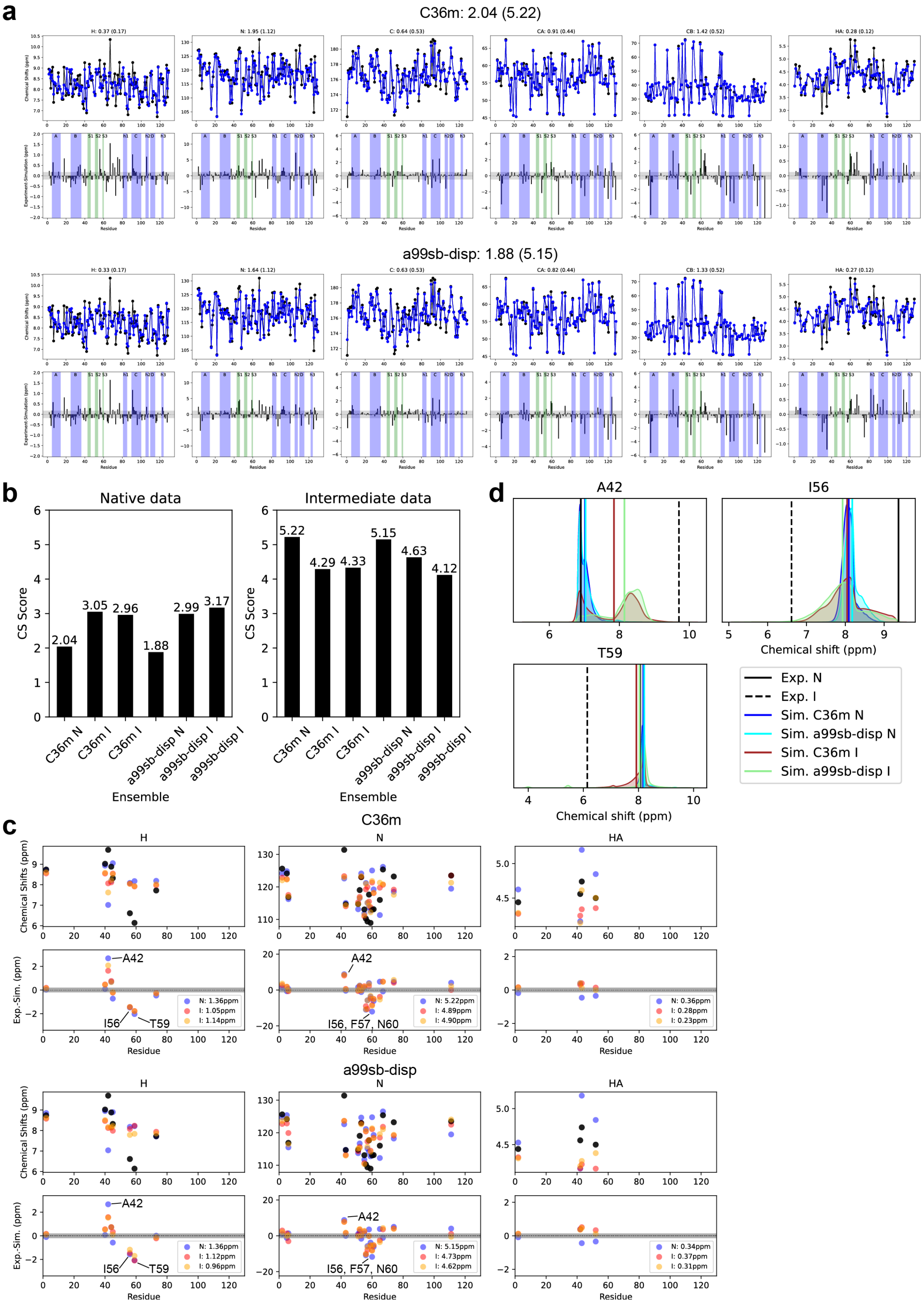


**Figure S10.** Analysis and comparison of back-calculated NMR chemical shifts with experiments. (a) Comparison of experimental the back-calculated backbone chemical shifts of the native state. The top row contains the experimental values (black) and ensemble-averaged values obtained by MD (4x2.5μs). The bottom row shows the difference between the experimental and simulated values. The shaded area in grey represents the average error of the forward model (see Methods). For each force field, the average chemical shift score (see Methods) for the native and intermediate (in parentheses) state are shown. (b) Agreement between all MD ensembles and the experimental data for the native (left) and intermediate (right) state, quantified by the chemical shift score. (c) Comparison of experimental the back-calculated backbone chemical shifts of the intermediate state. For each force field, the top row shows the experimental (black) and simulated values, and the bottom row shows the difference of the simulated values from the experimental data (grey shaded areas highlight the average uncertainty of the forward model). Residues 42, 56, 57, and 59 exhibit the largest deviation from the experimental values. The average deviation for each nucleus is shown in the figure legend of the bottom row. (d) Probability distributions of the sampled amide proton chemical shifts by MD in the native and intermediate states. The experimental values are indicated with vertical lines in black and MD ensemble average are also shown in vertical lines.
